# YAP1 Activation by Human Papillomavirus E7 Promotes Basal Cell Identity in Squamous Epithelia

**DOI:** 10.1101/2021.11.10.468068

**Authors:** Joshua Hatterschide, Paola Castagnino, Hee Won Kim, Steven M. Sperry, Kathleen T. Montone, Devraj Basu, Elizabeth A. White

**Affiliations:** Department of Otorhinolaryngology: Head and Neck Surgery, University of Pennsylvania Perelman School of Medicine, Philadelphia, PA, USA; Department of Otolaryngology-Head and Neck Surgery, Aurora St. Luke’s Medical Center, Milwaukee, Wisconsin, U.S.A.; Department of Pathology and Laboratory Medicine, University of Pennsylvania Perelman School of Medicine, Philadelphia, PA, USA

## Abstract

Persistent human papillomavirus (HPV) infection of stratified squamous epithelial cells causes nearly five percent of cancer cases worldwide. HPV-positive oropharyngeal cancers harbor few mutations in the Hippo signaling pathway compared to HPV-negative cancers at the same anatomical site, prompting the hypothesis that an HPV-encoded protein inactivates the Hippo pathway and activates the Hippo effector YAP1. The HPV E7 oncoprotein is required for HPV infection and for HPV-mediated oncogenic transformation. We investigated the effects of HPV oncoproteins on YAP1 and found that E7 activates YAP1, promoting YAP1 nuclear localization in basal epithelial cells. YAP1 activation by HPV E7 required that E7 bind and degrade the tumor suppressor PTPN14. E7 required YAP1 transcriptional activity to extend the lifespan of primary keratinocytes, indicating that YAP1 activation contributes to E7 carcinogenic activity. Maintaining infection in basal cells is critical for HPV persistence, and here we demonstrate that YAP1 activation causes HPV E7 expressing cells to be retained in the basal compartment of stratified epithelia. We propose that YAP1 activation resulting from PTPN14 inactivation is an essential, targetable activity of the HPV E7 oncoprotein relevant to HPV infection and carcinogenesis.

## Introduction

Human papillomaviruses (HPV) are non-enveloped viruses with circular double-stranded DNA genomes that infect keratinocytes in stratified squamous epithelia (Doorbar et al., 2015; Graham, 2017; McBride, 2017). Although most HPV infections are cleared by the immune system, some infections persist and form higher grade lesions that can lead to cancer (Koshiol et al., 2008; McBride, 2021; Radley et al., 2016; Rositch et al., 2013). HPV infection at mucosal epithelial sites causes cancers including oropharyngeal, cervical, vaginal, penile, and anal malignancies (de Martel et al., 2017; Gillison et al., 2015). Nearly 5% of human cancer cases are caused by persistent infection with one of the high-risk (oncogenic) human papillomavirus genotypes (de Martel et al., 2020).

Inactivation of host cell tumor suppressors by the high-risk HPV E6 and E7 oncoproteins modulates cellular processes that enable HPV persistence. Two well-characterized instances of tumor suppressor inactivation by HPV are high-risk HPV E6 proteins targeting p53 for proteasome-mediated degradation and high-risk HPV E7 proteins binding and degrading the retinoblastoma protein (RB1) (Heck et al., 1992; Münger et al., 1989; Scheffner et al., 1990; Seavey et al., 1999; Werness et al., 1990). Both p53 degradation and RB1 inactivation are required for productive HPV infection (Collins et al., 2005; Flores et al., 2000; Kho et al., 2013; McLaughlin-Drubin et al., 2005; Wang et al., 2009). In addition to supporting productive infection, E7 is essential for HPV-mediated carcinogenesis (Mirabello et al., 2017). The impact of the HPV oncoproteins on cell growth control pathways is reflected in human cancer genomic data: genes in the p53 pathway and in the RB1-related cell cycle pathway are frequently mutated in HPV-negative head and neck squamous cell carcinoma (HNSCC) but infrequently mutated in HPV-positive HNSCC (Sanchez-Vega et al., 2018).

Although some of the growth-promoting activities of high-risk HPV E6 and E7 are well established, open questions remain. RB1 binding/degradation by high-risk HPV E7 is necessary but insufficient for E7 transforming activity (Balsitis et al., 2006, 2005; Banks et al., 1990; Ciccolini et al., 1994; Helt and Galloway, 2002; Huh et al., 2005; Ibaraki et al., 1993; Jewers et al., 1992; Phelps et al., 1992; Strati and Lambert, 2007; White et al., 2015). Papillomavirus researchers have sought to identify one or more activities of HPV E7 that cooperate with RB1 inactivation to promote carcinogenesis and to identify the cellular pathway affected by such an activity. Human cancer genomic data indicates that like the p53 and cell cycle pathways, the Hippo signaling pathway is more frequently mutated in HPV-negative than in HPV-positive HNSCC. The core Hippo pathway consists of a kinase cascade upstream of the effector proteins Yes-Associated Protein (YAP1) and its paralogue TAZ. When the Hippo kinases are inactive, YAP1 and TAZ are activated and translocate to the nucleus. In stratified squamous epithelia YAP1 is primarily expressed in the basal layer, where YAP1 activation is regulated by contextual cues including cell density, tension in the extracellular matrix, and contact with the basement membrane (Elbediwy et al., 2016; Totaro et al., 2017; Zhang et al., 2011). In normal stratified squamous epithelia, activation of YAP1 and TAZ promotes expansion of the basal cell compartment, and inhibition of YAP1 and TAZ allows keratinocytes to differentiate (Beverdam et al., 2013; Elbediwy and Thompson, 2018; Schlegelmilch et al., 2011; Totaro et al., 2017; Yuan et al., 2020; Zhang et al., 2011). Mutations in many of the tumor suppressors upstream of YAP1/TAZ are common in a variety of cancer types (Moroishi et al., 2015).

Non-receptor protein tyrosine phosphatase 14 (PTPN14) has been implicated as a tumor suppressor and negative regulator of YAP1 (Knight et al., 2018; Mello et al., 2017; Poernbacher et al., 2012; Wang et al., 2012). Diverse HPV E7 bind directly to PTPN14 and recruit the E3 ligase UBR4 to direct PTPN14 for proteasome-mediated degradation (Szalmás et al., 2017; White et al., 2016, 2012b; Yun et al., 2019). We have shown that PTPN14 degradation and RB1 binding/degradation are separable activities of HPV E7 that each contribute to E7 carcinogenic activity (Hatterschide et al., 2020, 2019; White et al., 2016). However, the downstream consequences of PTPN14 degradation are poorly understood, and so far we have not observed that PTPN14 inactivation in human keratinocytes causes an increase in canonical YAP1 target genes *CTGF* and *CYR61*.

These observations regarding an additional transforming activity of HPV E7, the ability of E7 to inactivate PTPN14, and the relative paucity of mutations in the Hippo pathway in HPV-positive HNSCC led us to hypothesize that HPV E7-mediated activation of YAP1 is required for the transforming activity of high-risk HPV E7. Here we show that expression of high-risk HPV E7 is sufficient to activate YAP1 and that HPV E7 requires YAP1/TAZ-TEAD transcriptional activity to promote cell growth. We demonstrate that HPV E7 must bind PTPN14 to activate YAP1 and that PTPN14 inactivation alone is sufficient to activate YAP1. YAP1 activation by HPV E7 is restricted to the basal layer of the epithelium where we found *PTPN14* expression to be enriched. Our finding that either HPV E7 or PTPN14 loss activate YAP1 specifically in basal epithelial cells led us to investigate the role of YAP1 activation during normal HPV infection. HPV infection begins in basal epithelial keratinocytes (Day and Schelhaas, 2014; Pyeon et al., 2009; Roberts et al., 2007) and infected basal cells are the site of persistent HPV infection. The basal cell compartment contains the only long-lived cells in the epithelium and the HPV genome can be maintained in dividing basal cells without productive replication (Egawa et al., 2012; Parish et al., 2006; You et al., 2004). Activation of YAP1 and TAZ has been proposed to maintain the progenitor cell state in several different epithelia (Beverdam et al., 2013; Heng et al., 2020; Hicks-Berthet et al., 2021; Szymaniak et al., 2015; Yimlamai et al., 2014; Zhao et al., 2014). If YAP1 activation by E7 promotes the maintenance of a basal cell state in stratified squamous epithelia, YAP1 activation could facilitate the persistence of HPV-positive cells. Testing this hypothesis, we found that YAP1 activation and PTPN14 degradation by E7 both promote the maintenance of cells in the basal compartment of stratified epithelia. We propose that YAP1 activation facilitates HPV persistence and contributes to the carcinogenic activity of high-risk HPV E7.

## Results

### HPV E7 activates YAP1 in basal keratinocytes

A comprehensive analysis of somatic mutations and copy number variations in human tumor samples revealed that the cell cycle, p53, and Hippo pathways are the three pathways that exhibit the greatest difference in alteration frequency in HPV-negative vs HPV-positive HNSCC (Sanchez-Vega et al., 2018). We used data made available by The Cancer Genome Atlas (TCGA) through cBioPortal (Lawrence et al., 2015) to recapitulate the finding that genes in these pathways are altered at a lower frequency in HPV-positive than in HPV-negative HNSCC (Figure 1A and Figure 1—figure supplement 1). However, most HPV-positive HNSCC arise in the oropharynx. We repeated the analysis of pathway alteration rates using data only from HPV-positive and HPV-negative oropharyngeal squamous cell carcinomas (OPSCC) (Figure 1A and Figure 1—figure supplement 1). Consistent with previous findings, HPV-negative OPSCC were more frequently altered in the p53, cell cycle, and Hippo pathways than HPV-positive OPSCC. Many of the Hippo pathway alterations in HPV-negative HNSCC or OPSCC are amplification of the YAP1/TAZ oncogenes or inactivating mutation in an upstream inhibitor of YAP1/TAZ. Either alteration type is consistent with a carcinogenic role for YAP1 activation in HNSCC.

**Figure 1.**
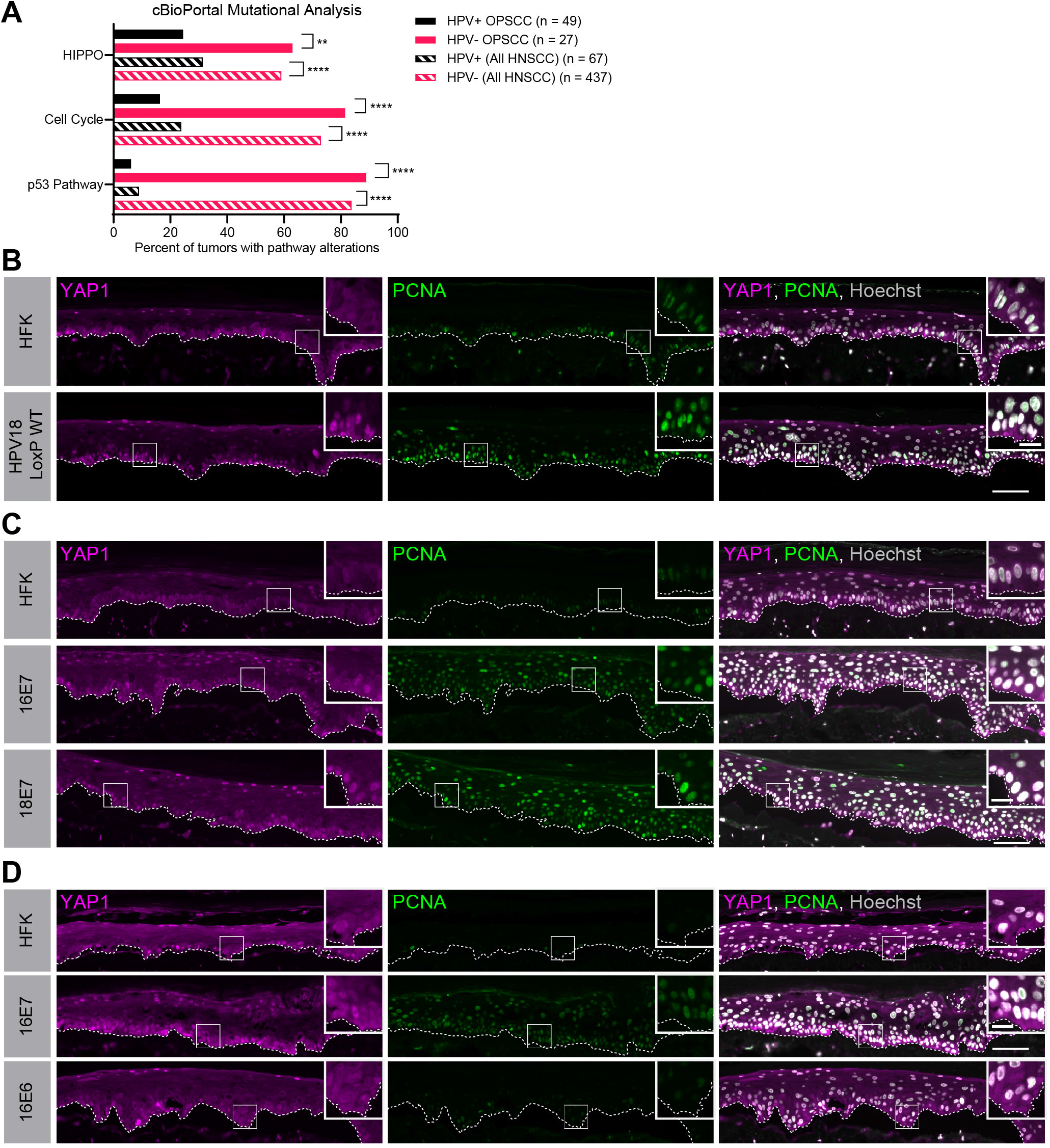
HPV E7 activates YAP1 in basal epithelial keratinocytes. (A) cBioPortal analysis for total genomic mutations and copy number alterations in HPV+/- OPSCC and HNSCC. Graph displays the percent of tumors with alterations in each pathway. Statistical significance was determined by Fisher’s exact test. (B-D) Organotypic cultures were grown from primary HFK, HFK harboring the HPV18 genome, or HFK transduced with retroviral expression encoding HPV E6 or E7 proteins. FFPE sections of cultures grown from (C) HFK or HFK harboring the HPV18 genome, (D) HFK or HFK expressing HPV16 E7 or HPV18 E7, or (E) HFK or HFK expressing HPV16 E6 or HPV16 E7 were stained for YAP1 (magenta), PCNA (green), and Hoechst (gray). White dashed lines indicate the basement membrane. White boxes indicate the location of insets in main images. Main image scale bars = 100 μm. Inset scale bars = 25 μm.

To test whether an HPV-encoded protein activates YAP1, we grew three dimensional (3D) organotypic epithelial cultures to model the differentiation of keratinocytes into basal and suprabasal compartments. Organotypic cultures of primary human foreskin keratinocytes (HFK) harboring an HPV18 genome exhibited increased YAP1 staining and increased YAP1 nuclear localization, indicative of YAP1 activation, particularly in the basal layer of the epithelium, compared to HFK cultures (Figure 1B and Figure 1—figure supplement 2A,B). Proliferating cell nuclear antigen (PCNA) transcription increases upon RB1 inactivation and is a marker of HPV E7 expression. In contrast to the basal layer-specific compartmentalization of YAP1 activation in the HPV18 genome containing cells, PCNA levels were increased in these cultures in both the basal and suprabasal layers of the epithelium.

We next tested whether high-risk HPV E6 or E7 alone was sufficient to activate YAP1. HFK transduced with retroviral expression vectors encoding HPV16 E6, HPV16 E7, or HPV18 E7 were used to grow organotypic cultures. YAP1 expression and nuclear localization were increased in the HPV16 E7 and HPV18 E7 expressing cells relative to parental HFK cells (Figure 1C and Figure 1—figure supplement 3A-C). As in the HPV18 genome-containing cells, YAP1 activation was restricted to the basal epithelial layer. YAP1 expression or nuclear localization did not increase in organotypic cultures of HPV16 E6 expressing cells (Figure 1D and Figure 1— figure supplement 4). Constitutive expression of either HPV16 E7 or HPV18 E7 induced PCNA expression in basal and suprabasal cells. We conclude that HPV promotes increased YAP1 expression and nuclear localization in basal keratinocytes and that E7 is sufficient for YAP1 activation.

### HPV E7 activates YAP1 in keratinocytes through PTPN14 degradation

We previously discovered that HPV E7 targets the YAP1 inhibitor PTPN14 for proteasome-mediated degradation (White et al., 2016, 2012b). We tested whether loss of PTPN14 expression in keratinocytes was sufficient to activate YAP1 in stratified epithelia by growing 3D organotypic cultures from previously described control and PTPN14 knockout (KO) N/Tert-Cas9 keratinocytes (Hatterschide et al., 2019). We found that YAP1 levels and YAP1 nuclear localization were increased in PTPN14 KO cultures compared to controls (Figure 2A and Figure 2—figure supplement 1A-C). YAP1 activation in basal epithelial cells lacking PTPN14 was comparable to YAP1 activation in HPV E7 cells. We conclude that loss of PTPN14 expression activates YAP1 in basal keratinocytes.

**Figure 2.**
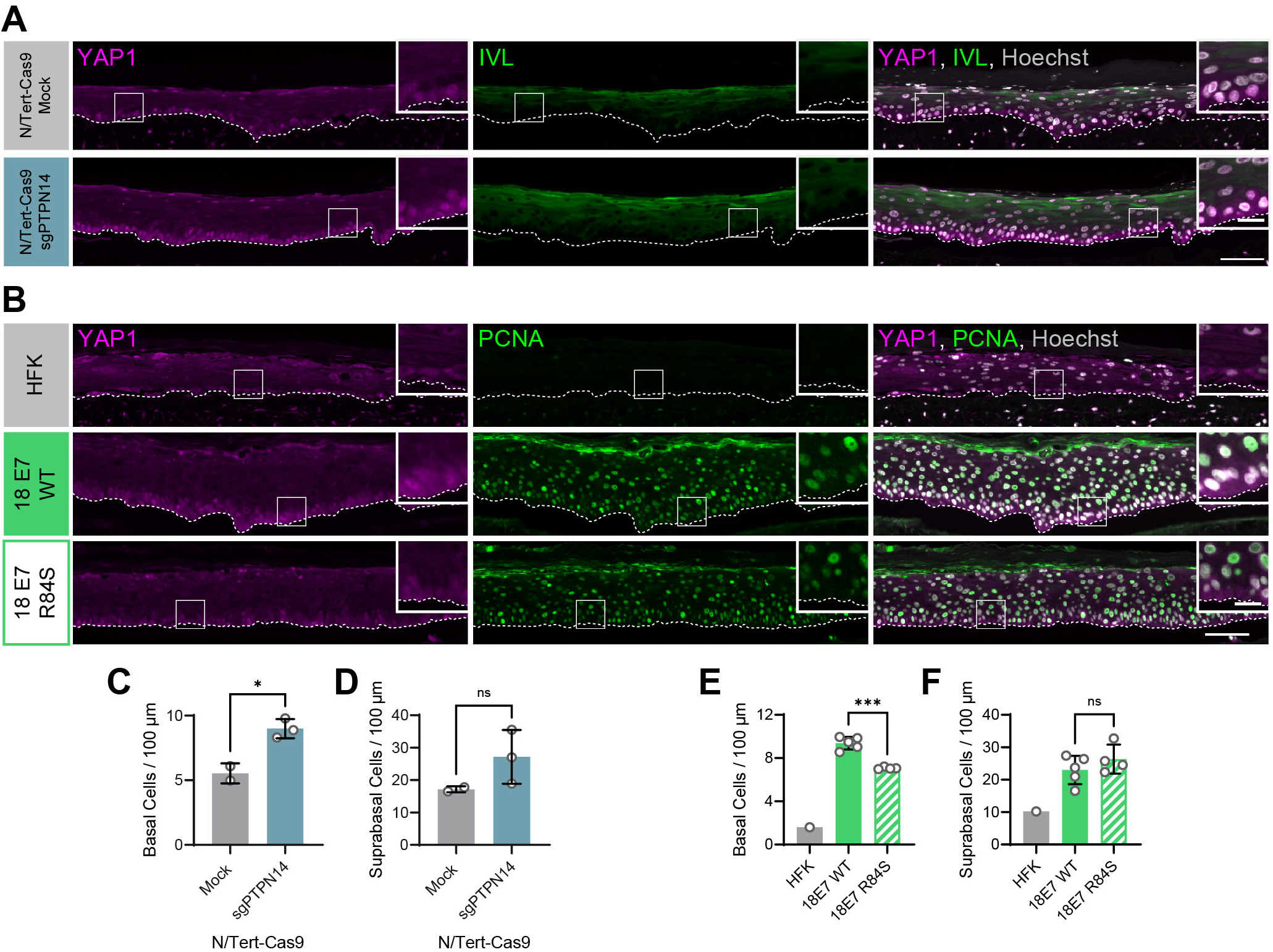
HPV E7 activates YAP1 in basal keratinocytes through PTPN14 degradation. Organotypic cultures were grown from N/Tert-Cas9 keratinocytes or primary HFK transduced with retroviral expression vectors encoding HPV18 E7 WT or R84S. (A) FFPE sections of cultures grown from mock or sgPTPN14 transfected N/Tert-Cas9 keratinocytes were stained for YAP1 (magenta), IVL (green), and Hoechst (Gray). (B) FFPE sections of cultures grown from parental HFK, HPV18 E7 WT or HPV18 E7 R84S expressing HFK were stained for YAP1 (magenta), PCNA (green), and Hoechst (Gray). White dashed lines indicate the basement membrane. White boxes indicate the location of insets in main images. Main image scale bars = 100 μm. Inset scale bars = 25 μm. (C-F) Quantification of the number of (C and E) basal cells and (D and F) suprabasal cells per 100 μm of epidermis. Graphs display the mean ± SD and each individual data point (independent cultures). Statistical significance was determined by ANOVA (*p<0.05, ***p<0.001).

A highly conserved C-terminal arginine in E7 makes a direct interaction with the C-terminus of PTPN14, and the HPV18 E7 R84S variant is unable to bind or degrade PTPN14 (Hatterschide et al., 2020; Yun et al., 2019). To test whether PTPN14 degradation by HPV E7 is required for activation of YAP1, we grew 3D organotypic cultures using primary HFK transduced with retroviral expression vectors encoding HPV18 E7 wild type (WT) or HPV18 E7 R84S. Indeed, YAP1 expression and nuclear localization in the basal layer of HPV18 E7 R84S cultures were reduced compared to HPV18 E7 WT controls (Figure 2B and Figure 2—figure supplement 2).

In addition to activating YAP1, PTPN14 loss increased basal cell density from an average of 5.5 cells per 100 μm in control cultures to 9.0 cells per 100 μm in PTPN14 KO cultures (Figure 2C). Basal cell density was higher in HPV18 E7 WT cultures (9.4 cells per 100 μm) than in HPV18 E7 R84S cultures (to 7.1 cells per 100 μm) (Figure 2E). No statistically significant difference in suprabasal cell density was observed in either comparison (Figure 2D,F). We conclude that E7 expression or PTPN14 loss in stratified squamous epithelia is sufficient to activate YAP1 in the basal layer of the epithelium and increase basal cell density.

### PTPN14 expression is enriched in basal keratinocytes

YAP1 activation was restricted to basal epithelial cells in our organotypic cultures leading us to hypothesize that PTPN14 may act as a basal layer specific inhibitor of YAP1. We therefore sought to determine whether *PTPN14* expression is restricted to a specific subset of cells in the stratified epithelium. In a recent single cell-RNA seq analysis of human neonatal foreskin epidermis, *PTPN14* mRNA expression was enriched in the basal-III cluster, a subset of basal cells predicted to differentiate directly into spinous cells (Figure 3A,B) (S. Wang et al., 2020). *PTPN14* expression was higher in basal-III cells than in the spinous or granular cell clusters. To test whether *PTPN14* expression is higher in basal or suprabasal cells in our cultures, we used laser capture microdissection to isolate basal and suprabasal layers from 3D organotypic cultures grown from unmodified primary HFK (Figure 3C). We found that there was a ∼5-fold enrichment of *PTPN14* mRNA in the basal epithelial layer compared to the suprabasal layers (Figure 3D). As expected, the basal integrins *ITGA6* and *ITGB4* were expressed in the basal layer (Figure 3E) and the differentiation markers *KRT1* and *IVL* were expressed in the suprabasal layers (Figure 3F). The same pattern of *PTPN14* mRNA expression was observed in an organotypic culture grown from primary HFK expressing HPV18 E7 WT (Figure 3—figure supplement 1A-C). We conclude that *PTPN14* mRNA is enriched in basal keratinocytes in the presence or absence of HPV E7. Our data support that PTPN14 acts as a YAP1 inhibitor specifically in the basal compartment of stratified epithelia.

**Figure 3.**
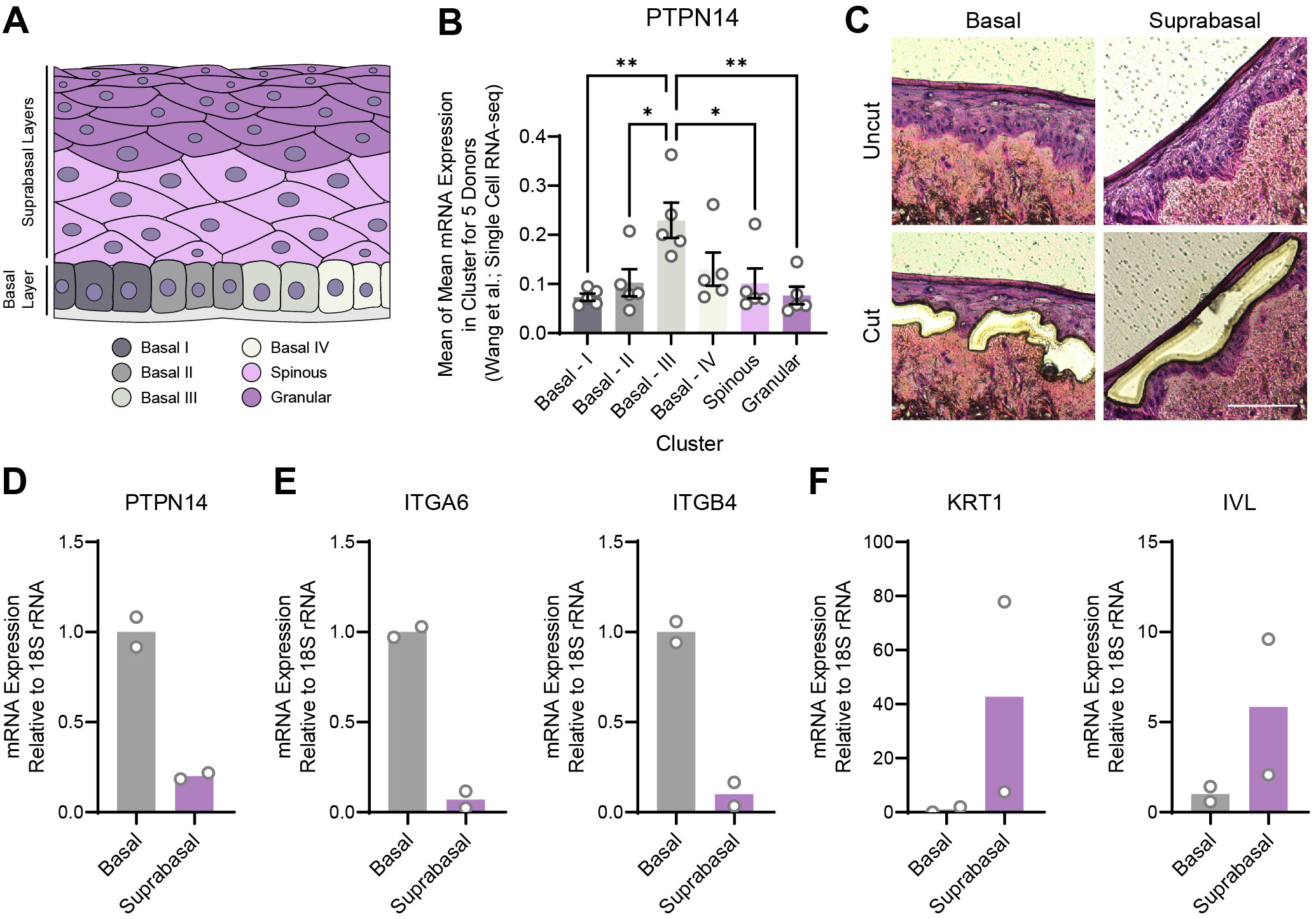
PTPN14 expression is enriched in basal keratinocytes. (A-B) Single-cell RNA sequencing data and clustering analysis from Wang et al. was reanalyzed to assess PTPN14 expression in different subsets of epidermal cells. (A) Diagram of epidermis; shading depicts tissue localization of cell clusters. (B) For each donor, the mean of PTPN14 mRNA expression was calculated for each cell cluster. Graphs display the mean of PTPN14 mRNA expression for each donor (circles) as well as the mean of all five donors ± SEM (bars and error bars). Statistical significance was determined by ANOVA (*p<0.05, **p<0.01). (C-F) Basal and suprabasal layers from organotypic cultures were dissected using laser capture microdissection. (C) Representative images of HFK cultures before and after individual laser dissections. Hundreds of such cuts were performed per sample. (D-F) RNA was purified from isolated layers and qRT-PCR was used to assess the expression of PTPN14 (D), basal cell markers ITGA6 and ITGB4 (E), and differentiation markers KRT1 and IVL (F). Graphs display the mean and each individual data point.

### YAP1/TAZ regulate differentiation downstream of PTPN14

In previous unbiased experiments we found that the primary effect of PTPN14 inactivation on transcription is to repress epithelial differentiation gene expression (Hatterschide et al., 2020, 2019). However, we also observed that PTPN14 inactivation did not increase expression of the canonical YAP1/TAZ targets *CTGF* and *CYR61*. Consistent with this difference there was minimal overlap between PTPN14-dependent differentially expressed genes and the genes listed in the MSigDB conserved YAP1 signature (Figure 4A). We therefore asked whether the ability of PTPN14 to regulate differentiation gene expression requires YAP1/TAZ as intermediates. Transduction of keratinocytes with a PTPN14 lentivirus induced the expression of the differentiation markers *KRT10* and *IVL* in a dose-dependent manner (Figure 4—figure supplement 1A-C). To test whether PTPN14 required YAP1/TAZ to increase *KRT1* and *IVL*, we transfected HFK with siRNAs targeting *YAP1* and *WWTR1* then transduced the cells with PTPN14 lentivirus (Figure 4B). HFK transfected with control siRNA exhibited the expected increase in *KRT1* and *IVL* after transduction with PTPN14 lentivirus (Figure 4C,D and Figure 4—figure supplement 2A,B). However, keratinocytes depleted of YAP1/TAZ did not express relatively more *KRT1* or *IVL* when PTPN14 was overexpressed than when it was not. We conclude that PTPN14 requires YAP1 and/or TAZ to regulate differentiation gene expression in keratinocytes. Both pairs of YAP1/TAZ siRNA had the same effect on differentiation in response to PTPN14 overexpression yet only one pair efficiently depleted TAZ protein levels (Figure 4B), leading us to speculate that YAP1 is the key intermediate connecting PTPN14 levels to differentiation gene expression.

**Figure 4.**
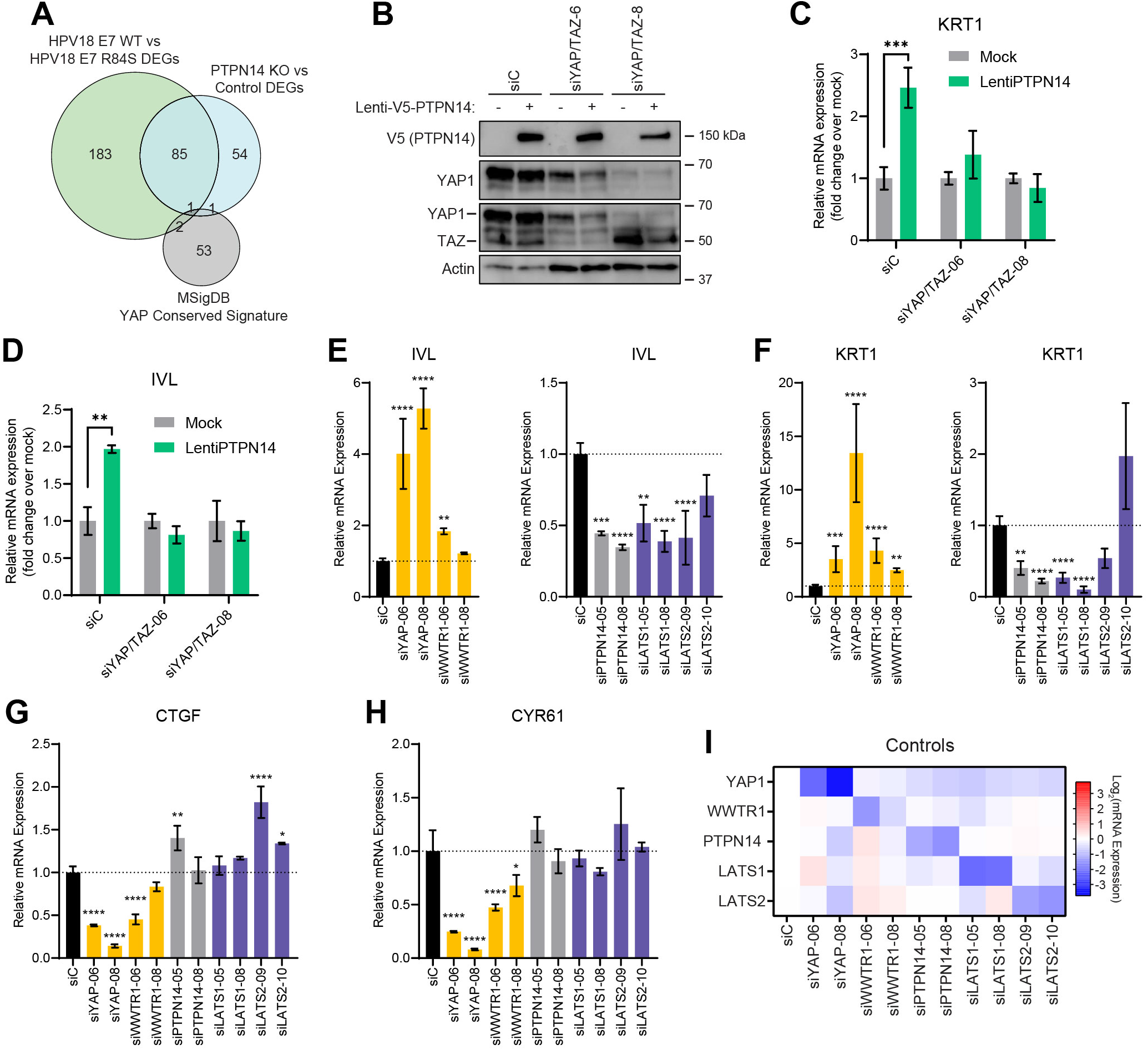
YAP1/TAZ regulate differentiation downstream of PTPN14. (A) Venn diagram comparing the MSigDB YAP conserved signature to the differentially expressed genes (DEG) from our two published experiments that reflect PTPN14 loss in keratinocytes. (B-D) *YAP1* and *WWTR1* were simultaneously knocked down by siRNA transfection in HFK. Transfected HFK were then transduced with PTPN14 lentivirus at 24h post transfection. Cells were lysed for protein and total cellular RNA at 72h post transfection. (B) Cell lysates were subjected to SDS/PAGE/Western analysis and probed with antibodies to PTPN14, YAP1, TAZ, and Actin. (C and D) qRT-PCR was used to measure the expression of the differentiation markers *KRT1* and *IVL* relative to *G6PD*. Graphs display fold change in gene expression relative to the mock transduced cells. (E-I) Primary HFK were transfected with siRNAs targeting *YAP1*, *WWTR1* (TAZ), *PTPN14*, *LATS1*, and *LATS2*. Two siRNAs were used per target. qRT-PCR was used to measure gene expression for: the differentiation markers *IVL* (E) and *KRT1* (F), and the canonical YAP1/TAZ targets *CTGF* (G) and *CYR61* (H). Data confirming that individual siRNA transfections depleted intended transcripts is summarized in a heatmap of log_2_(fold-change) levels (I). Bar graphs display the mean ± SD of three independent replicates. Statistical significance was determined by ANOVA (*p<0.05, **p<0.01, ***p<0.001, ****p<0.0001).

Next, we tested whether repression of keratinocyte differentiation occurs upon loss of LATS1 and LATS2, the core Hippo pathway kinases that phosphorylate and inhibit YAP1 and TAZ. We used siRNAs to deplete *PTPN14*, *LATS1*, or *LATS2* and measured the expression of the differentiation markers *KRT1* and *IVL* (Figure 4E,F). Depletion of *PTPN14, LATS1,* or *LATS2* all decreased differentiation gene expression to a similar degree. Consistent with our previous experiments, none of the three knockdowns significantly affected the levels of *CTGF* or *CYR61* (Figure 4G-H). Direct depletion of *YAP1* or *WWTR1* affected both differentiation gene expression and *CTGF/CYR61* levels. *YAP1* knockdown always had a stronger effect than did *WWTR1* knockdown and our qRT-PCR analyses supported that *WWTR1* transcript levels were low in HFK. This result shows that inactivation of three different YAP1 inhibitors dampens differentiation gene expression and does not increase canonical YAP1 target gene expression in keratinocytes. Taken together, these data support that PTPN14 promotes differentiation through inhibition of YAP1/TAZ despite not affecting canonical YAP1/TAZ target genes.

### HPV-positive HNSCC are less differentiated than HPV-negative HNSCC

We next asked whether the gene expression pattern observed downstream of PTPN14 loss is reflected in HPV-positive cancers. HPV-positive HNSCC have a strong propensity toward poorly differentiated, basaloid histology (Mendelsohn et al., 2010; Pai and Westra, 2009), which is reflected in their transcriptional profile (Hatterschide et al., 2019). We confirmed the relationship between HPV positivity and greater impairment of differentiation by immunohistochemical analysis of the differentiation marker KRT1 in sections of 14 HPV-negative tumors and 48 HPV-positive tumors (Figure 5A). 43% of HPV-negative tumors and 12.5% of HPV-positive tumors stained positive for KRT1. We additionally measured gene expression in patient-derived xenograft (PDX) models generated from human HNSCC. We measured *KRT1*, *KRT10*, and *IVL* levels using RNA extracted from 11 HPV-negative and 8 HPV-positive HNSCC PDX. Each differentiation marker was expressed at a markedly lower level in HPV-positive PDX than in HPV-negative PDX (Figure 5B). We observed the same pattern of differentiation marker gene expression in an analysis of transcriptomic data from other cohorts (Figure 5—figure supplement 1A-C) (Lawrence et al., 2015). Having confirmed that HPV-positive HNSCC exhibit reduced expression of differentiation markers than do HPV-negative HNSCC, we measured *CTGF* and *CYR61* levels. We found no significant difference in expression of these canonical YAP1/TAZ target genes in HPV-positive vs HPV-negative PDX, although there was a trend towards higher *CTGF* in the HPV-positive PDX (Figure 5C and Figure 5—figure supplement 1D,E). The pattern of low expression of differentiation markers and unchanged canonical YAP1/TAZ target gene expression in HPV-positive versus HPV-negative patient samples is consistent with the effects of PTPN14 inactivation in cultured cells.

**Figure 5.**
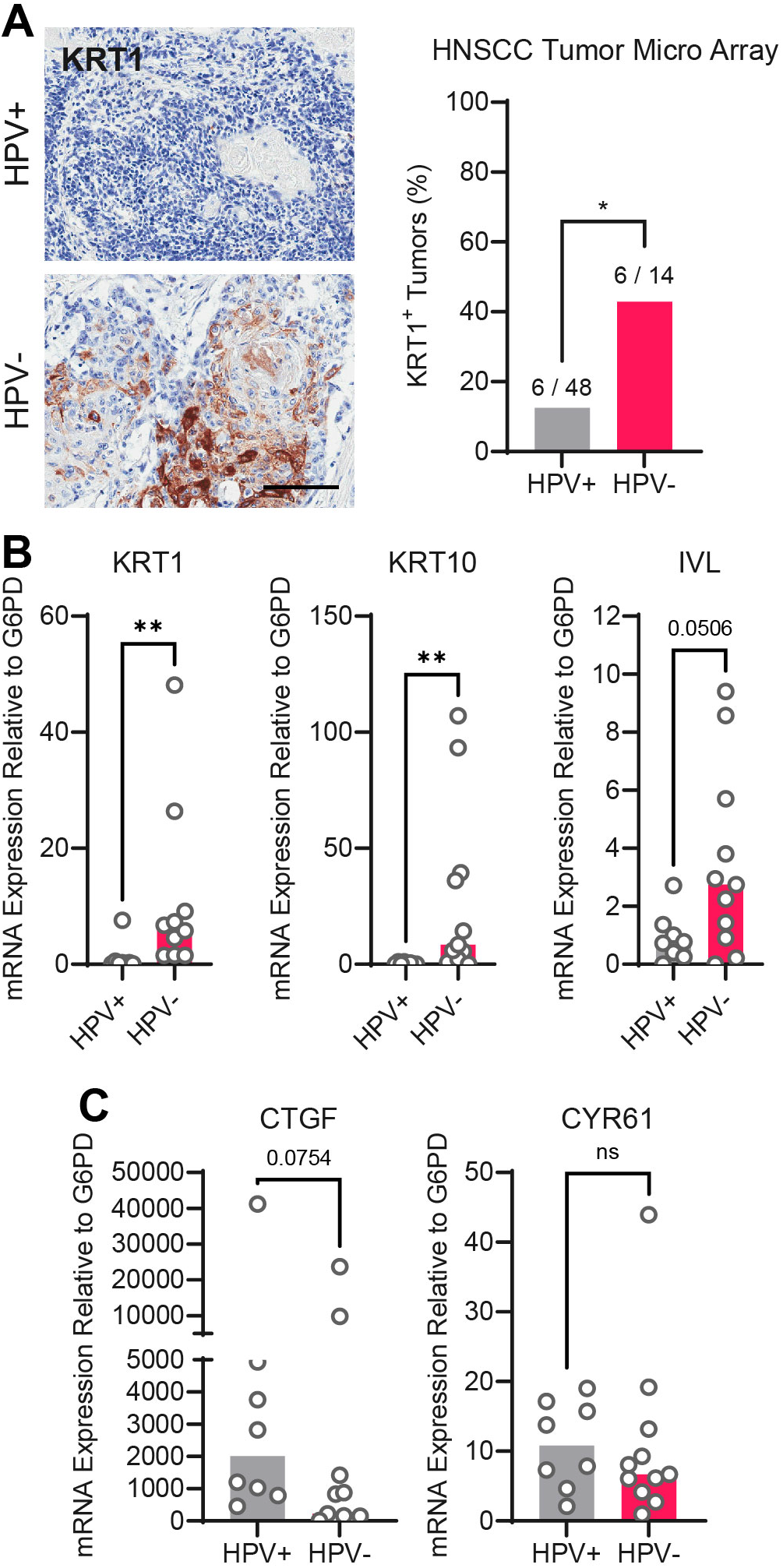
HPV-positive HNSCC are less differentiated than HPV-negative HNSCC. (A) Human HNSCC tumor samples were stained for KRT1 (left). Scale bar = 100 μm. Graph displays the percentage of tumors that were KRT1^+^ (right). Statistical significance was determined by Fisher’s exact test. (B-C) Total RNA was purified from PDX samples and qRT-PCR was used to assess gene expression of (B) the differentiation markers KRT1, KRT10, and IVL and (C) the canonical YAP1/TAZ targets CTGF and CYR61. Statistical significance was determined by Mann-Whitney nonparametric test. (*p<0.05, **p<0.01, ****p<0.0001).

### High-risk HPV E7 require YAP1/TAZ-TEAD transcriptional activity to extend the lifespan of primary keratinocytes

High-risk but not low-risk HPV E7 proteins can extend the lifespan of primary keratinocytes (Halbert et al., 1991). The TEADi protein is a genetically encoded competitive inhibitor that prevents binding between YAP1/TAZ and TEAD transcription factors (Yuan et al., 2020). We used TEADi to test whether YAP1/TAZ-TEAD transcriptional activity was required for high-risk HPV E7 to extend the lifespan of primary HFK. We transduced HFK with retroviral vectors encoding GFP, HPV16 E7, or HPV18 E7 plus a lentiviral vector encoding doxycycline-inducible GFP-TEADi. As anticipated, HPV16 E7 or HPV18 E7 extended the lifespan of primary HFK based on cumulative population doublings (Figures 6A,B). TEADi induction upon doxycycline treatment decreased the lifespan of primary HFK in the presence or absence of E7, but the effect of YAP1/TAZ-TEAD inhibition was greater in the HPV16 E7 and HPV18 E7 cells, where E7 had minimal ability to promote growth in the presence of TEADi. We conclude that high-risk HPV E7 proteins require YAP1/TAZ-TEAD transcriptional activity for their lifespan extending capacity in primary keratinocytes.

**Figure 6.**
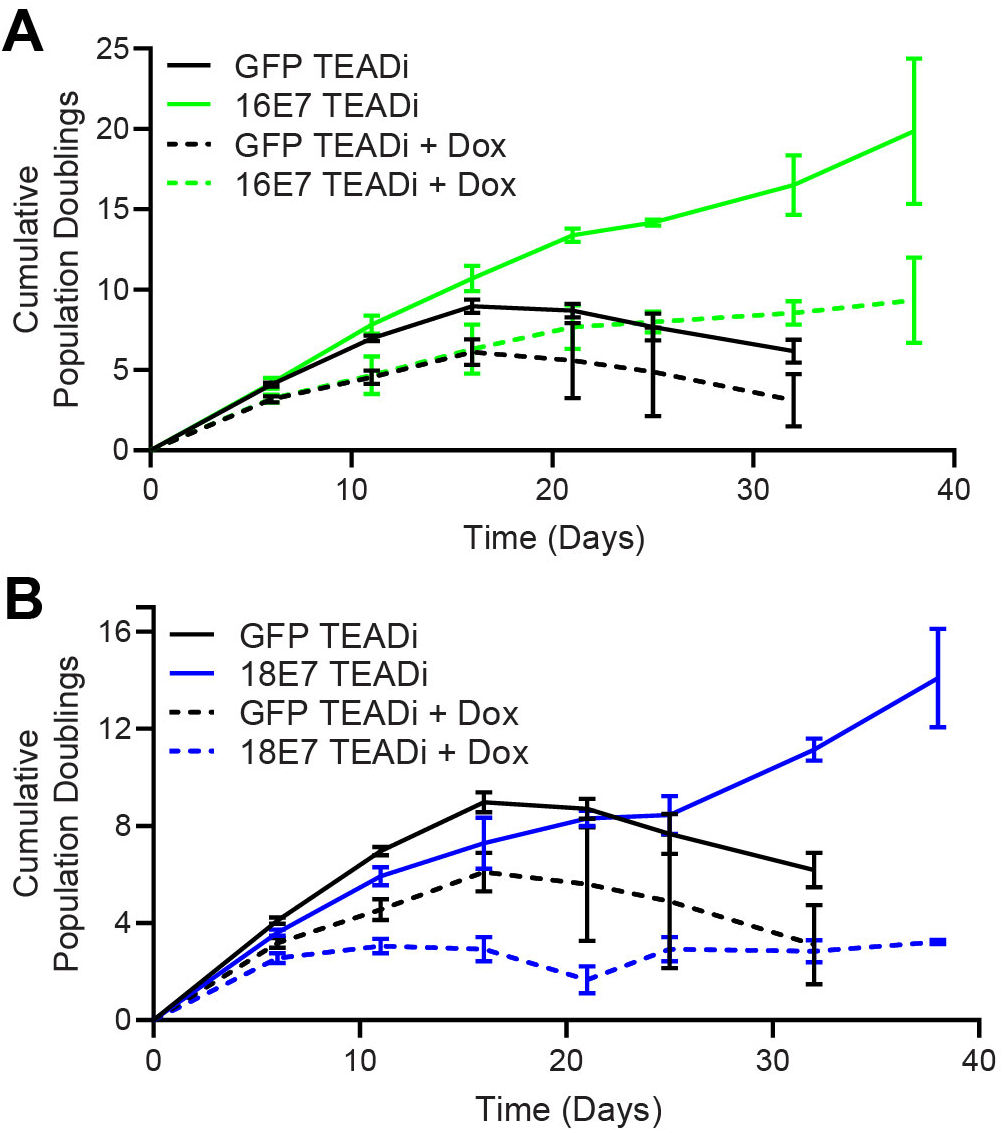
High-risk HPV E7 requires YAP1/TAZ-TEAD transcriptional activity to extend the lifespan of primary keratinocytes. Primary HFK were transduced with retroviruses encoding HPV16 E7, HPV18 E7, or GFP, plus pInducer20 TEADi lentivirus. Each cell population was cultured with or without 1 μg/mL doxycycline in the media for 38 days and population doublings were tracked with each passage. Graph displays the mean ± SD of two independently transduced cell populations per condition.

### PTPN14 loss and YAP1 activation promote basal cell retention in organotypic cultures

YAP1 overexpression impairs differentiation and promotes progenitor cell identity in squamous and non-squamous epithelia. HPV infection is maintained in a reservoir of infected basal cells and productive virus replication begins upon commitment to differentiation. To better understand how repression of differentiation downstream of YAP1 activation affects HPV viral biology, we developed an assay to measure cell retention in the basal epithelial layer. We hypothesized that YAP1 activation by HPV E7 might promote the adoption of a basal cell identity in stratified squamous epithelia. In our cell fate monitoring assay, a small proportion of GFP-labeled cells were mixed with unmodified, parental HFK, and the pool was used to generate organotypic cultures in which normal labeled cells are randomly distributed throughout the epithelium.

Our initial experiment tested whether YAP1 activation altered cell fate in stratified squamous epithelia. We used GFP-labeled tracing cells that expressed doxycycline-inducible YAP1 WT, YAP1 S127A (hyperactive), or YAP1 S94A (cannot bind TEAD transcription factors) (Figure 7—figure supplement 1A,B). In organotypic cultures grown from a 1:25 mixture of GFP-labeled cells and unmodified HFK, about 20% of uninduced GFP+ cells were found in the basal layer. Induction of YAP1 WT or YAP1 S127A expression was sufficient to promote the retention of nearly 60% of labeled cells in the basal layer of the epithelium (Figure 7A,B). Only around 40% of GFP+ cells were found in the basal layer when YAP1 S94A was induced. These data indicate that YAP1 activation causes cells to be retained in the basal layer of a stratified squamous epithelium. The ability of YAP1 to bind TEAD transcription factors contributed to its activity in the cell fate assay.

**Figure 7.**
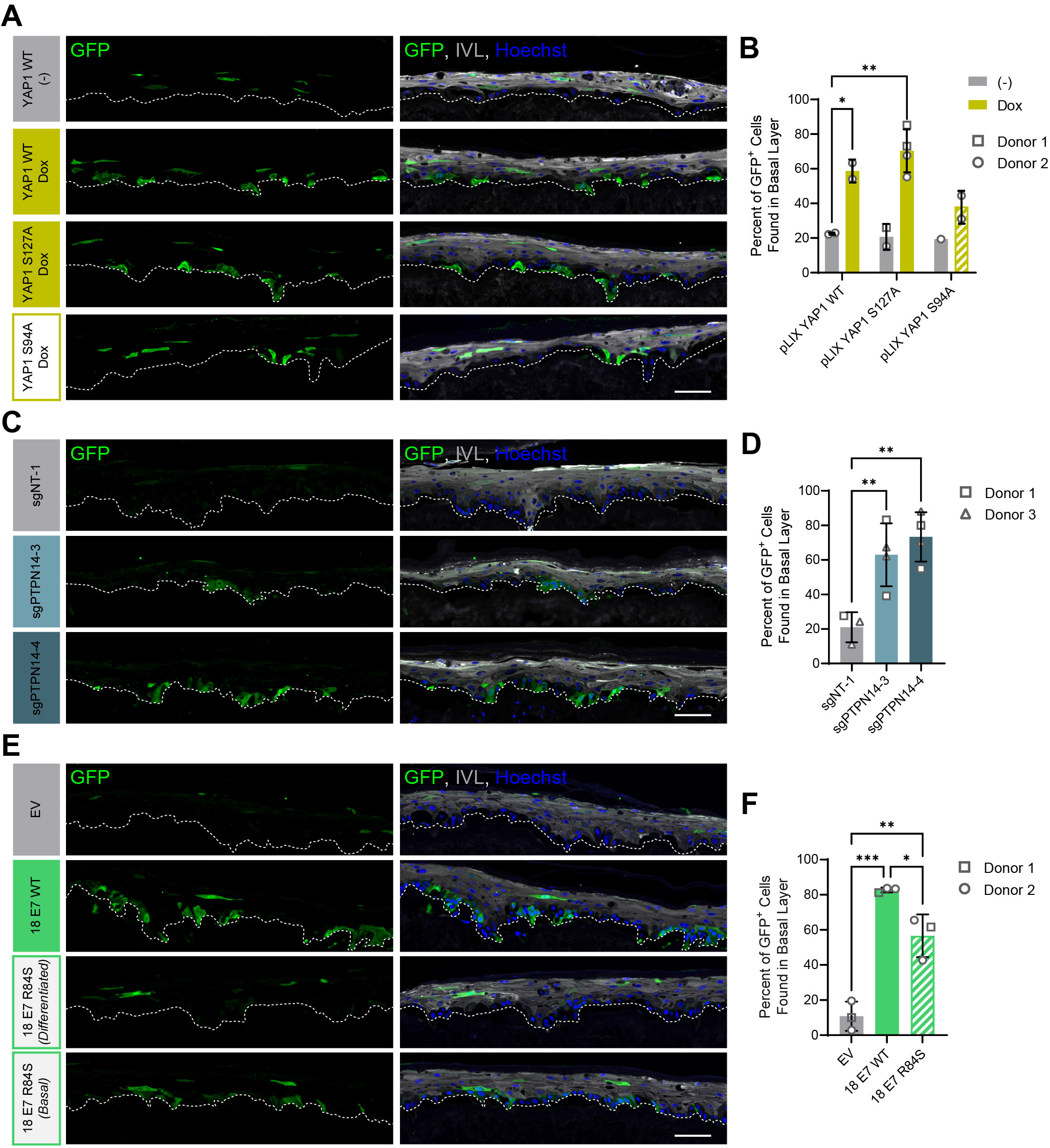
PTPN14 loss and YAP1 activation by HPV E7 promote basal cell retention in organotypic cultures. Organotypic cultures were grown from GFP-labeled HFK mixed with unmodified HFK. (A-B) GFP-labeled HFK were transduced with lentiviral vectors encoding YAP1 WT, YAP1 S127A, or YAP1 S94A under the control of a doxycycline inducible promoter. GFP-labeled YAP1 cells were mixed 1:25 into unmodified HFK and organotypic cultures were grown from the mixture. Cultures were grown +/- 1 μg/mL doxycycline. (C-D) GFP-labeled HFK were transduced with LentiCRISPR v2 vectors encoding control or PTPN14 targeting sgRNAs. GFP-labeled cells were mixed 1:25 into unmodified HFK and organotypic cultures were grown from the mixture. (E-F) GFP-labeled HFK were transduced with HPV18 E7 WT, HPV18 E7 R84S, or the empty vector (EV). GFP-labeled HPV18 E7 cells were mixed 1:50 into unmodified HFK and organotypic cultures were grown from the mixture. (A, C, E) FFPE sections of cultures were stained for GFP (green), IVL (grey), and Hoechst (blue). Scale bar = 100 μm. (B, D, F) Quantification of the percentage of GFP+ cells found in the basal layer. Graphs display the mean ± SD and each individual data point (independent cultures). Shapes indicate cultures grown from different HFK donors. Statistical significance was determined by ANOVA. (*p<0.05, **p<0.01).

We next tested whether loss of PTPN14 expression was sufficient to promote basal cell identity. We grew organotypic cultures from mixtures of unmodified primary HFK and GFP-labeled control or PTPN14 KO HFK (Figure 7—figure supplement 1C,D). 60-70% of PTPN14 KO tracer cells were found in the basal layer when either of two PTPN14 guide RNAs were used whereas about 20% of control tracer cells were retained in the basal layer (Figure 7C,D). Thus, PTPN14 knockout is sufficient to promote basal cell fate determination in keratinocytes.

Next, we tested whether HPV E7 promoted basal cell retention and if so, whether its cell retention activity required PTPN14 degradation. We grew organotypic cultures from mixtures of GFP-labeled HFK expressing HPV18 E7 WT, HPV18 E7 R84S, or the empty vector control diluted 1:50 into unmodified primary HFK (Figure 7—figure supplement 1E,F). We found that nearly 80% of GFP-labeled HPV18 E7 WT tracer cells were retained in the basal layer compared to about 10% of labeled control cells (Figure 7E,F). HPV18 E7 WT labeled cells were numerous and grouped in clusters in the basal layer, suggesting that E7 promoted the clonal expansion of labeled basal cells. Both effects were dampened in experiments using HPV18 E7 R84S tracer cells (cannot degrade PTPN14). Labeled HPV18 E7 R84S cells exhibited varying degrees of basal cell expansion and basal cell retention and approximately 60% of labeled cells were in the basal layer. HPV18 E7 R84S retains the ability to inactivate RB1 and we interpret these data to mean that the proliferation of labeled basal cells resulted from RB1 inactivation. Finally, HPV18 E7 ΔDLLC cannot bind RB1 but can bind and degrade PTPN14. In a cell fate experiment using GFP-labeled HPV18 E7 ΔDLLC tracer cells, the labeled cells were present mainly as single cells in the basal layer (Figure 7—figure supplement 2A-B). The behavior of the two mutant HPV E7 proteins supports that PTPN14 degradation is required for basal cell retention and RB1 inactivation is required for basal cell expansion. We conclude that PTPN14 degradation and YAP1 activation by HPV18 E7 promote basal cell retention.

## Discussion

YAP1 and TAZ are oncogenes that promote growth and inhibit differentiation in stratified squamous epithelia (Elbediwy et al., 2016; Schlegelmilch et al., 2011; Totaro et al., 2017; Yuan et al., 2020; Zhang et al., 2011). Here we report that HPV E7 activates YAP1 (Figure 1). YAP1/TAZ-TEAD transcriptional activity is required for the carcinogenic activity of HPV E7 (Figure 6) and YAP1 activation by E7 biases HPV E7-expressing cells to be retained in the basal epithelial layer (Figure 7). Based on these findings we propose that YAP1 activation by HPV E7 enables HPV-infected cells to persist in stratified epithelia. There is substantial evidence that RB1 inactivation is necessary but insufficient for the transforming activity of high-risk HPV E7 (Balsitis et al., 2006, 2005; Banks et al., 1990; Ciccolini et al., 1994; Helt and Galloway, 2002; Huh et al., 2005; Ibaraki et al., 1993; Jewers et al., 1992; Phelps et al., 1992; Strati and Lambert, 2007; White et al., 2015). We propose that YAP1 activation cooperates with RB1 inactivation to enable the transforming activity of HPV E7.

PTPN14 binding by HPV18 E7 was required for activation of YAP1 in the basal layer and PTPN14 KO was sufficient for the same effect (Figure 2). Highly conserved amino acids in E7 participate in binding to PTPN14 (Hatterschide et al., 2020; Yun et al., 2019), indicating that YAP1 activation and maintenance of basal cell state is likely shared among diverse papillomavirus E7 proteins. Some minor genotype-specific differences were apparent. HPV18 E7 depletes PTPN14 protein levels more efficiently than HPV16 E7 (Hatterschide et al., 2020; White et al., 2016), which is consistent with the observed stronger effect of HPV18 E7 on YAP1 nuclear localization in basal cells (Figure 1). Genotype-specific differences could also explain the stronger effect of TEADi on HPV18 E7 in lifespan extension assays (Figure 6). Although other reports have suggested that HPV might activate YAP1 (He et al., 2015; Morgan et al., 2020; Olmedo-Nieva et al., 2020; Webb Strickland et al., 2018), no specific activity of an HPV protein has previously been shown to enable YAP1 activation. Other groups have proposed that HPV E6 activates YAP1 (He et al., 2015; Webb Strickland et al., 2018), but we did not observe YAP1 activation by HPV E6. We conclude that activation of YAP1 by HPV E7 is contingent upon its ability to bind and degrade PTPN14.

Even when HPV E7 was expressed in all layers of a stratified epithelium, YAP1 levels and nuclear localization increased only in basal epithelial cells. We found that E7 required PTPN14 degradation to activate YAP1 and that PTPN14 was expressed predominantly in basal keratinocytes (Figure 3). Basal cell-specific expression of *PTPN14* is consistent with the observation that it is regulated by p63, the master regulator of basal cell identity in stratified epithelia (Perez et al., 2007). We propose that PTPN14 inhibits YAP1 primarily in basal cells and that unlike the effects of E7 on RB1 in both differentiated and undifferentiated cells, E7 activates YAP1 primarily in basal cells.

Degradation of PTPN14 by HPV E7 represses keratinocyte differentiation but does not induce canonical Hippo pathway target genes (Hatterschide et al., 2020, 2019). Nonetheless, we found that PTPN14 overexpression promoted differentiation only in the presence of YAP1/TAZ (Figure 4C,D). Few studies have tested how YAP1 inhibitor inactivation alters gene expression downstream of YAP1. Here we demonstrate that inactivation of LATS1 or LATS2, two well-characterized inhibitors of YAP1/TAZ, also repressed differentiation genes but did not induce canonical YAP1/TAZ targets (Figure 4E-I). Taken together, these experiments indicate that PTPN14 acts through YAP1/TAZ to regulate differentiation in keratinocytes. It is so far unclear why *CTGF* and *CYR61* expression is sensitive to large changes in total levels of YAP1 or TAZ yet is unaffected by alterations in regulators upstream of YAP1/TAZ. Nonetheless, the pattern of low differentiation gene expression and unchanged expression of canonical YAP1/TAZ target genes caused by PTPN14 loss is consistent with gene expression differences between HPV-positive and HPV-negative HNSCC.

PTPN14 knockout and knockdown reduced differentiation gene expression in monolayer culture. Even so, we did not observe reduced differentiation in suprabasal layers of organotypic cultures grown from PTPN14 knockout cells (Figure 2A and Figure 2—figure supplement 1A-C). Using our cell fate monitoring assay, we determined that instead, HPV18 E7 promotes basal cell retention and that either YAP1 overexpression or PTPN14 KO are sufficient for this activity (Figure 7). The effect of YAP1 activation on cell fate in our assay resembles several experiments in which YAP1 promotes progenitor cell identity in airway and liver epithelia (Yimlamai et al., 2014; Zhao et al., 2014). Our findings demonstrate that YAP1 activation enables basal cell fate determination in stratified squamous epithelia and show that loss of an inhibitor of YAP1 has the same effect. We conclude that one consequence of YAP1 activation by HPV E7 is that E7-expressing cells are retained in the basal layer of stratified squamous epithelia.

Although persistent infection is a prerequisite for HPV-mediated carcinogenesis, the mechanisms used by papillomaviruses to establish persistent infections remain incompletely understood. Maintaining infection in the basal cell compartment is critical for papillomavirus persistence. Substantial effort has been devoted to the mechanistic understanding of how the papillomavirus genome is stably maintained in the basal layer upon cell division. However, much less is known about how papillomaviruses manipulate epithelial cell fate to establish and expand the pool of infected basal cells. Previously, HPV E7 was believed to be primarily required to establish a cellular environment conducive to HPV DNA replication in suprabasal cells. We propose that a so far unappreciated role of E7 is that it activates YAP1 to facilitate HPV persistence by biasing infected cells to remain in the basal layer of the epithelium. Not every HPV E7-expressing cell was retained in the basal layer, so we do not anticipate that YAP1 activation would block differentiation-dependent HPV replication. HPV E6 also represses differentiation gene expression in keratinocytes and has been proposed to promote basal cell retention (Kranjec et al., 2017). Further research is needed to determine the extent to which different HPV genotypes depend on the activities of E6 or E7 for basal cell retention activity.

To the best of our knowledge, no other viruses are recognized to modulate cell fate decisions in solid tissues in a way that facilitates persistence. Some herpesviruses impact the choice between progenitor/differentiated cell fates in infected immune cells, for example Epstein-Barr Virus (EBV) restricts B-cell differentiation to facilitate viral latency (Knox and Carrigan, 1992; Niiya et al., 2006; Onnis et al., 2012; Romeo et al., 2019; Styles et al., 2017). Herpesviruses, polyomaviruses, and hepadnaviruses encode proteins proposed to activate YAP1/TAZ or alter Hippo signaling (Hwang et al., 2014; Liu et al., 2014, 2015; Nguyen et al., 2014; Shanzer et al., 2015; Tian et al., 2004; Z. Wang et al., 2020). Not all of the mechanisms used by these viruses to activate YAP1 nor the downstream consequences of YAP1 activation have been well defined. Our finding that HPV E7 activates YAP1 to manipulate cell fate opens up an exciting new line of inquiry into how YAP1, TAZ, and the Hippo signaling pathway could impact viral infections by regulating tissue developmental processes.

YAP1 activation and PTPN14 are relevant to both viral and non-viral cancers. We found that a genetically encoded inhibitor of YAP1/TAZ-TEAD transcription inhibited the growth of high-risk HPV E7 expressing cells (Figure 6), indicating that high-risk HPV E7 proteins require YAP1 or TAZ for carcinogenesis. YAP1/TAZ activation is sufficient to drive carcinogenesis in mouse models of cervical and oral cancer (He et al., 2019; Nishio et al., 2020; Omori et al., 2020), and the YAP1 inhibitor verteporfin reduced the growth of HPV-positive tumors in a xenograft model (Liu et al., 2019). YAP1 activation correlates with the clinical stage of HPV infection (Nishio et al., 2020), and YAP1 localizes to the nucleus in HPV-positive cancers (Alzahrani et al., 2017). Basal cell carcinoma (BCC) is the non-viral cancer that is most clearly linked to PTPN14. Germline inactivating mutations in *PTPN14* are associated with a 4- to 8-fold increase in risk of BCC by age 70 (Olafsdottir et al., 2021) and somatic mutations in *PTPN14* are frequent in BCC (Bonilla et al., 2016). YAP1/TAZ-TEAD transcriptional activity also restricts differentiation in BCC cells (Yuan et al., 2021). We propose that the specific association of PTPN14 with BCC is related to our observation that PTPN14 loss activates YAP1 in basal epithelial cells. YAP1 inhibition is of major clinical interest for several cancer types, and it is appealing to speculate that targeting YAP1 could treat persistent HPV infection and/or HPV-positive cancers.

## Materials and Methods

### Plasmids and cloning

pInducer20 EGFP-TEADi was a gift from Ramiro Iglesias-Bartolome (Addgene plasmid # 140145) (Yuan et al., 2020). pQCXIH-Myc-YAP (Addgene plasmid # 33091), pQCXIH-Flag-YAP-S127A (Addgene plasmid # 33092), and pQCXIH-Myc-YAP-S94A (Addgene plasmid # 33094) were gifts from Kun-Liang Guan (Zhao et al., 2007). Each YAP1 ORF was amplified by PCR from pQCXIH, cloned into pDONR223, and transferred into pLIX_402 lentiviral backbone using Gateway recombination. pLIX_402 was a gift from David Root (Addgene plasmid # 41394). pLenti CMV GFP Hygro (656-4) was a gift from Eric Campeau & Paul Kaufman (Addgene plasmid # 17446) (Campeau et al., 2009). PHAGE-P-CMVt N-HA GFP was previously described (Galligan et al., 2014). pNeo-loxP-HPV18 was the kind gift of Thomas Broker and Louise Chow (Wang et al., 2009). The ΔDLLC mutation was introduced into the pDONR HPV18 E7 vector using site-directed mutagenesis. HPV18 E7 ΔDLLC and GFP ORFs were cloned into MSCV-P C-FlagHA GAW or MSCV-Neo C-HA GAW destination vectors using Gateway recombination. The remaining MSCV-P C-FlagHA and MSCV-Neo C-HA HPV E6 and HPV E7 retroviral plasmids and pHAGE lentiviral plasmids have been previously described (Hatterschide et al., 2020; White et al., 2016, 2012a, 2012b). A complete list of all plasmids used in this study is in Supplemental File 1.

### Cell culture, retrovirus production, and lentivirus production

Deidentified primary human foreskin keratinocytes (HFK) and human foreskin fibroblasts (HFF) were provided by the University of Pennsylvania Skin Biology and Disease Resource-Based Center (SBDRC). N/Tert-1 cells are hTert-immortalized HFK (Dickson et al., 2000), and N/Tert-Cas9 mock and sgPTPN14-1 are N/Tert-1 cells further engineered to constitutively express Cas9 (Hatterschide et al., 2019). Keratinocytes for cell fate experiments were cultured in keratinocyte serum-free media (KSFM) (Life Technologies, Carlsbad, California) mixed 1:1 with Medium 154 (Thermo Fisher Scientific, Waltham, Massachusetts) with the human keratinocyte growth supplement (HKGS) (Thermo Fisher Scientific) (Duperret et al., 2015; Egolf et al., 2019). Keratinocytes for all other experiments were cultured as previously described (White et al., 2012a). HFF were cultured in Dulbecco’s Modified Eagle Medium (DMEM) (Thermo Fisher Scientific) supplemented with antibiotic and antimycotic. HFK harboring the HPV18 genome were previously described (Hatterschide et al., 2020), and were generated by transfecting cells with the pNeo-loxP-HPV18 vector (Wang et al., 2009) along with NLS-Cre and selecting with G418 to generate a stable population. Lentiviruses and retroviruses were produced in 293T or 293 Phoenix cells respectively as previously described (White et al., 2016). Stable keratinocyte populations were generated following transduction by selection with puromycin, G418, or hygromycin alone or in combination.

### Lifespan extension assay

Primary HFK were engineered and cultured as described in cell culture, retrovirus production, and lentivirus production. The growth of engineered HFK was monitored in culture for 38 days. Population doublings were calculated using the number of cells at the beginning and end of each passage.

### Organotypic epithelial culture

Devitalized human dermis was provided as deidentified material from the University of Pennsylvania SBDRC. Stands for organotypic epithelial cultures were printed using high temperature, autoclavable resin at the University of Pennsylvania Biotech Commons 3D-printing facility. Organotypic cultures were generated as previously described (Duperret et al., 2015; Egolf et al., 2019). Devitalized dermis was seeded with primary HFF on the dermal side at a density of 3 x 10^4^ cells per cm^2^ of culturing area and cultured for four days. Dermis and fibroblasts were then stretched across 3D-printed stands. The epidermal side of the dermis was seeded with unmodified or engineered keratinocytes at a density of 1 x 10^6^ cells per cm^2^. Organotypic cultures were cultured in E media (Fehrmann and Laimins, 2005) with the dermal layer maintained at the air-liquid interface starting on the day of seeding keratinocytes. Cultures were allowed to stratify for 12-14 days, then trimmed and fixed in 10% neutral buffer formalin for 24 hours. Tissues were embedded in paraffin and sectioned by the SBDRC Core A. A complete list of all organotypic cultures used in this study is in Supplemental File 2.

### siRNA transfection

Primary HFK were transfected with siRNAs using the Dharmafect 1 transfection reagent. All siRNA experiments were collected 72 h post transfection. Two siRNAs were used to target each gene in an experiment. The siRNAs used in this study were all purchased from Dharmacon (Lafayette, Colorado): nontargeting siRNA, siYAP1-06, siYAP1-08, siWWTR1-06, siWWTR1-08, siPTPN14-05, siPTPN14-08, siLATS1-05, siLATS1-08, siLATS2-09, siLATS2-10.

### Laser capture microdissection

Formalin-fixed paraffin-embedded (FFPE) organotypic cultures were sectioned onto polyethylene naphthalate (PEN) membrane glass slides by the SBDRC Core A. Laser capture microdissection was performed on a Leica LMD 7000 microscope. Hundreds of microdissections were made per sample amounting to ∼1.5 mm^2^ of total dissected area per sample. RNA was isolated using the RNeasy FFPE kit (Qiagen, Germantown, Maryland). RNA concentration was determined using Qubit RNA HS assay kit (Life Technologies).

### Patient derived xenografts

The PDXs were previously established from surgical resections of treatment-naive HPV-positive OPSCC as described (Facompre et al., 2020). Human tumors were engrafted subcutaneously in NSG mice and passaged at least twice before cryopreservation when they reached a volume of 0.5-1.0 cm^3^. Total tumor RNA was isolated using the QIAamp RNA Blood Mini Kit (Qiagen).

### Western blotting

Western blots were performed using Mini-PROTEAN (Bio-Rad Laboratories, Hercules, California) or Criterion (Bio-Rad) Tris/Glycine SDS-PAGE gels and transfers were performed onto polyvinylidene difluoride (PVDF). Membranes were blocked with 5% nonfat dried milk in Tris-buffered saline with 0.05% Tween 20 (TBST). Membranes were incubated with primary antibodies as specified in Supplemental File 1. Following TBST washes, membranes were incubated with horseradish peroxidase-coupled secondary antibodies and imaged using chemiluminescent substrate on an Amersham Imager 600 (GE Healthcare, Chicago, Illinois).

### qRT-PCR

Unless otherwise specified, total cellular RNA was isolated using the NucleoSpin RNA extraction kit (Macherey-Nagel/Takara, San Jose, California). cDNA was generated from bulk RNA with the high-capacity cDNA reverse transcription kit (Applied Biosystems, Waltham, Massachusetts). cDNAs were used as a template for qPCR using Fast SYBR green master mix (Applied Biosystems) and a QuantStudio 3 system (Thermo Fisher Scientific). 18S rRNA qRT-PCR primers were ordered from Integrated DNA Technologies (Integrated DNA Technologies, Inc., Coralville, Iowa): FWD, 5- CGCCGCTAGAGGTGAAATTCT; REV, 5- CGAACCTCCGACTTTCGTTCT (Roh et al., 2005). KiCqStart SYBR green primers for qRT-PCR (MilliporeSigma, St. Louis, Missouri) were used for the remaining genes assayed in this study: KRT1, KRT10, IVL, ITGB4, ITGA6, CYR61, CTGF, PTPN14, YAP1, WWTR1, LATS1, LATS2, G6PD, and GAPDH.

### Immunofluorescence, immunohistochemistry, and microscopy

FFPE sections were prepared for immunofluorescence by deparaffinization with xylene washes, rehydration through an ethanol gradient, and heat induced epitope retrieval (HIER). Tissue sections were blocked with PBS containing 1% bovine serum albumin, 10% normal goat serum, and 0.3% Triton X-100. Tissue sections were incubated with primary antibodies at 4°C overnight, washed with PBS with 0.05% Tween 20, and incubated with fluorescently labeled secondary antibodies and Hoechst 33342 at room temperature. Antibody dilutions and HIER conditions are specified in Supplemental File 1. Fluorescent micrographs were captured using an Olympus IX81 microscope. All fluorescent micrograph images within the same figure panels were captured using the same exposure time and batch processed using the same contrast settings.

The TMA was constructed from surgical resection specimens of 120 HNSCC that vary by TNM stage and HPV status (Supplemental File 3). Archival FFPE tumors of the oral cavity and oropharynx were identified retrospectively and oropharyngeal tumors were evaluated for HPV status as per College of American Pathologists criteria (Lewis et al., 2018) using IHC for p16. When present, lymph node metastases were included in association with the primary tumor of origin. All FFPE specimens were represented in the TMA by at least three tissue cores that incorporate both non-necrotic central tumor regions and invasive margins. Tumor materials and clinical data were accessed under University of Pennsylvania IRB protocol 417200. Staining for KRT1 was performed by the Clinical Services Laboratory in the University of Pennsylvania Department of Pathology and Laboratory Medicine. Antibody information can be found in Supplemental File 1. The KRT1 stained slides were reviewed with a standard light microscope, and evaluation was based on the presence or absence of staining in the cytoplasm of tumor cells.

### Bioinformatic analysis

Genomic mutation and copy number variation data as well as tumor RNA-seq gene expression data from TCGA (Lawrence et al., 2015) were analyzed using the cBioPortal.org graphical interface (Cerami et al., 2012; Gao et al., 2013). RNA-seq V2 RSEM (RNA-Seq by Expectation Maximization) normalized expression values for individual genes were downloaded directly from cBioPortal.org. OPSCC were distinguished from HNSCC by clinical annotation of primary tumor site and HPV-positive and HPV-negative status was assigned based on previously reported HPV transcript status (Chakravarthy et al., 2016). Genes included as a part of each pathway analysis are listed in Supplemental File 4. Missense, truncating, and splice mutations of unknown significance as well as amplifications of tumor suppressor genes and deletion of oncogenes were excluded from total alteration tallies.

Single cell-RNA sequencing dataset derived from the human neonatal foreskin epidermis and subsequent clustering analysis were retrieved from GitHub (S. Wang et al., 2020) and reanalyzed with MATLAB. PTPN14 expression was calculated by averaging mRNA expression for all cells by cluster and donor.

## Acknowledgments

We thank the members of our laboratories, particularly Pavithra Rajagopalan, for helpful discussions. We thank Stephen M. Prouty, Ph.D. from the SBDRC for help with tissue processing and sectioning. Stands for organotypic cultures were printed courtesy of the University of Pennsylvania Libraries’ Biotech Commons. This work was supported by American Cancer Society grant 131661-RSG-18-048-01-MPC and NIH/NIAID R01 AI148431 to EAW. DB is supported by NIH/NIDCR R01 DE027185. JH was supported by NIH/NIAID T32 AI007324 and NIH/NIDCR F31 DE030365. SBDRC was funded by NIH grant P30 AR068589.

## Author Contributions

Conception and design: JH, EAW. Acquisition of data: JH, PC, HWK, KTM, EAW. Analysis and interpretation of data: JH, PC, KTM, DB, EAW. Drafting or revising the article: JH, PC, DB, EAW. Contributing unpublished essential data or reagents: SMS, KTM, DB.

---

**Figure 1—figure supplement 1.**
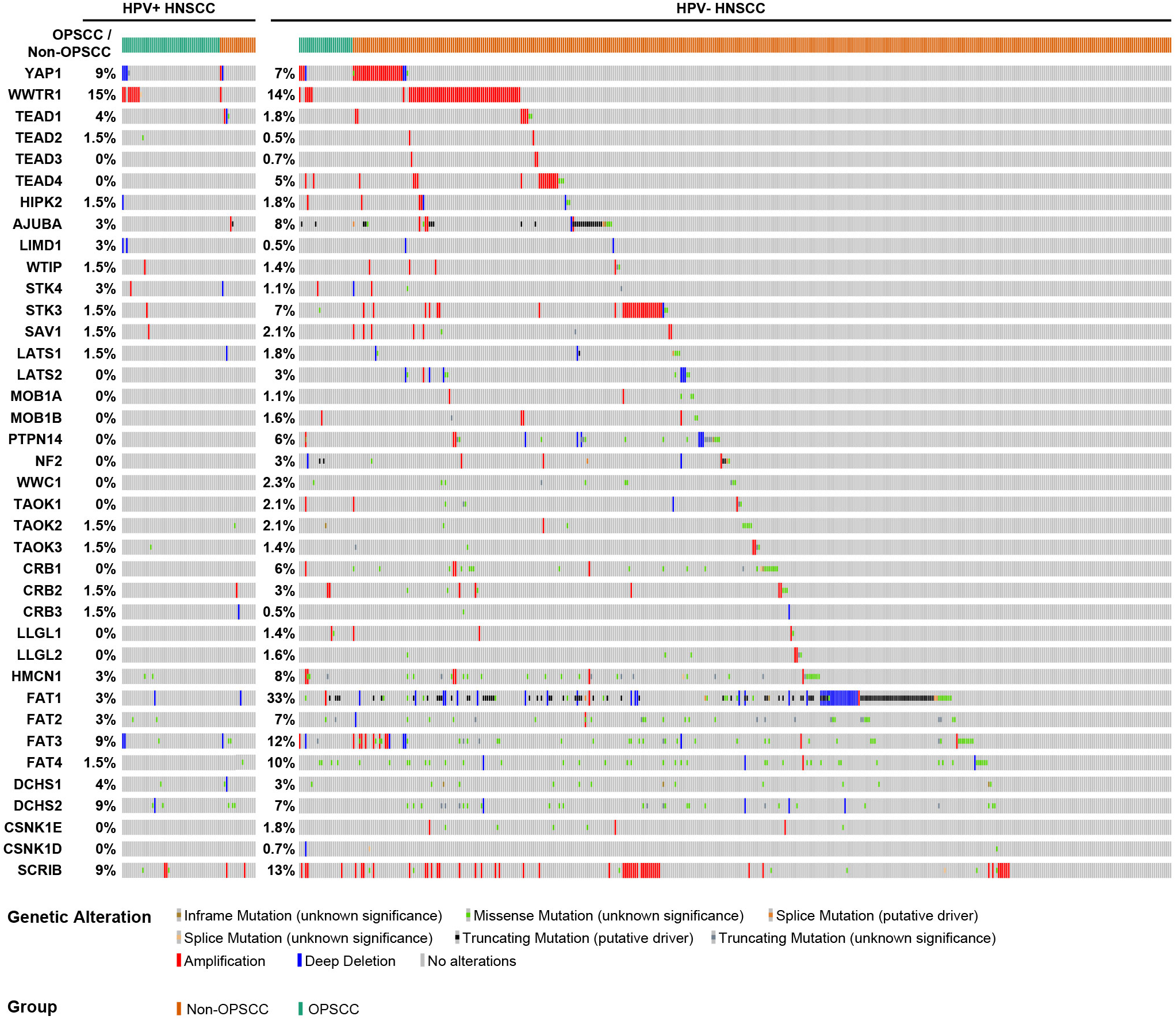
HPV-positive HNSCC have fewer Hippo pathway alterations and lower expression of differentiation genes. cBioPortal analysis for genomic mutations and copy number alterations in HPV+/- HNSCC and OPSCC. Oncoprint displays specific genomic alterations in individual tumor samples.

**Figure 1—figure supplement 2.**
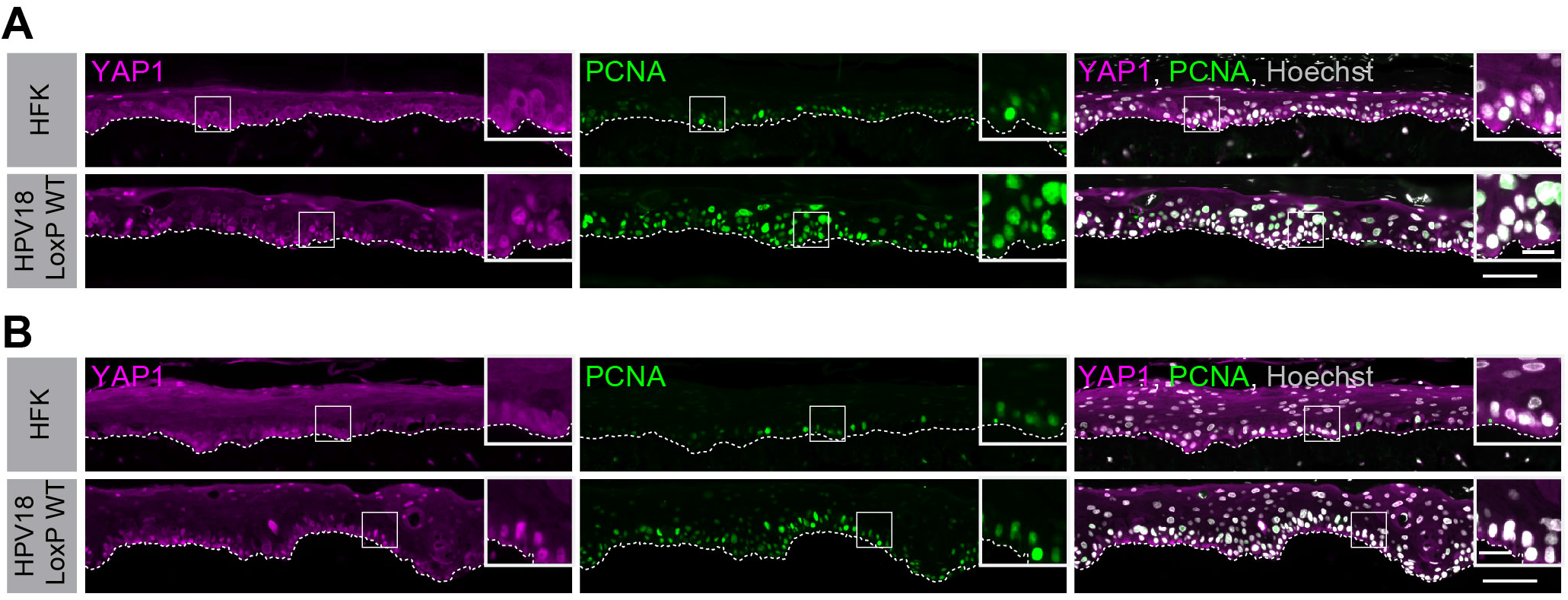
HPV18 E7 activates YAP1 in basal keratinocytes. (A-B) Additional replicates of organotypic cultures grown from primary HFK or HFK harboring the HPV18 genome. FFPE sections were stained for YAP1 (magenta), PCNA (green), and Hoechst (gray). White dashed lines indicate the basement membrane. White boxes indicate the location of insets in main images. Main image scale bars = 100 μm. Inset scale bars = 25 μm.

**Figure 1—figure supplement 3.**
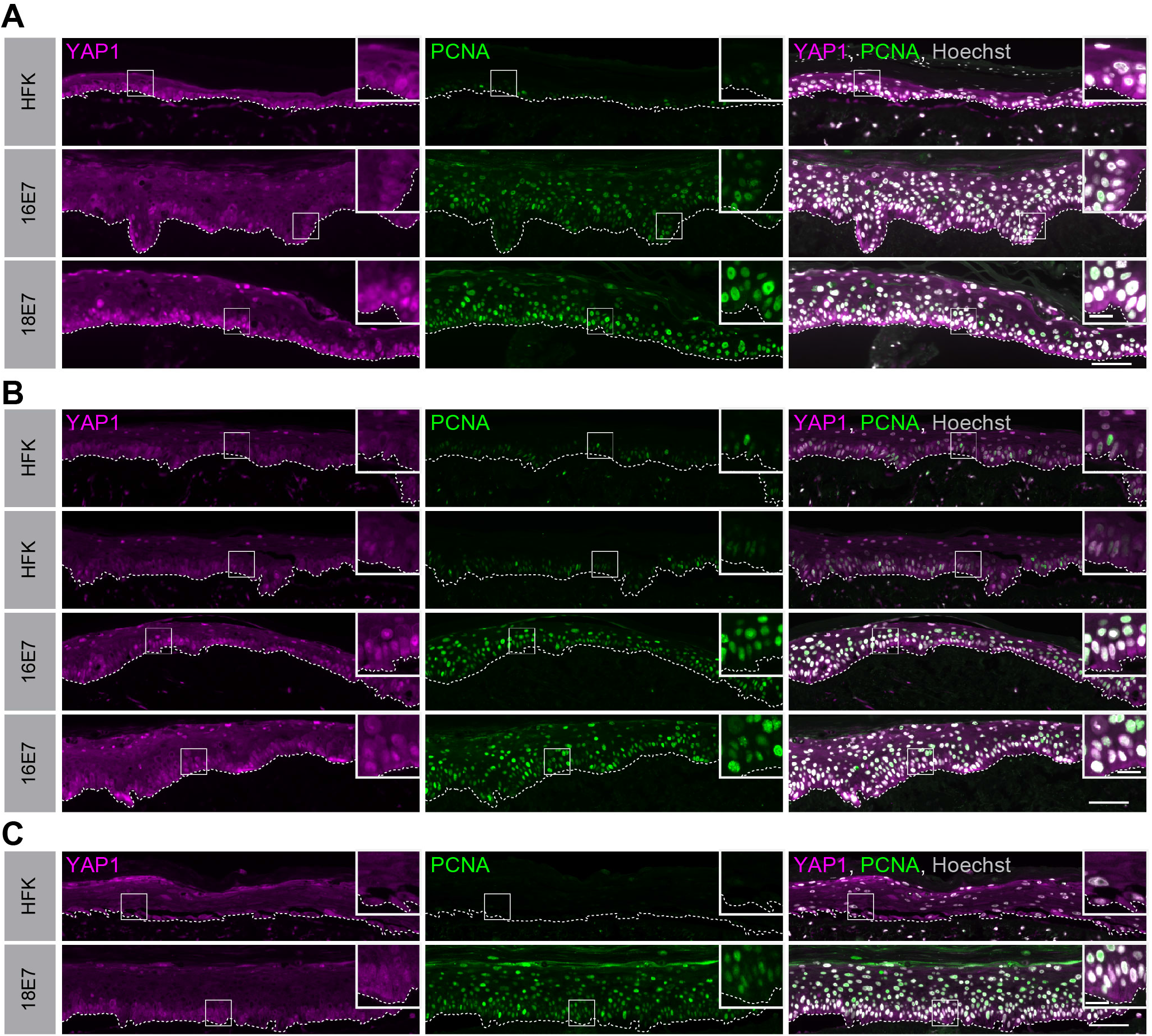
HPV E7 activates YAP1 in basal keratinocytes. Additional replicates of organotypic cultures grown from primary HFK or HFK transduced with retroviral expression encoding HPV E7 proteins. FFPE sections of cultures grown from (A) HFK or HFK expressing HPV16 E7 or HPV18 E7, (B) HFK or HFK transduced with HPV16 E7, or (E) HFK and HFK expressing HPV18 E7 were stained for YAP1 (magenta), PCNA (green), and Hoechst (gray). White dashed lines indicate the basement membrane. White boxes indicate the location of insets in main images. Main image scale bars = 100 μm. Inset scale bars = 25 μm.

**Figure 1—figure supplement 4.**
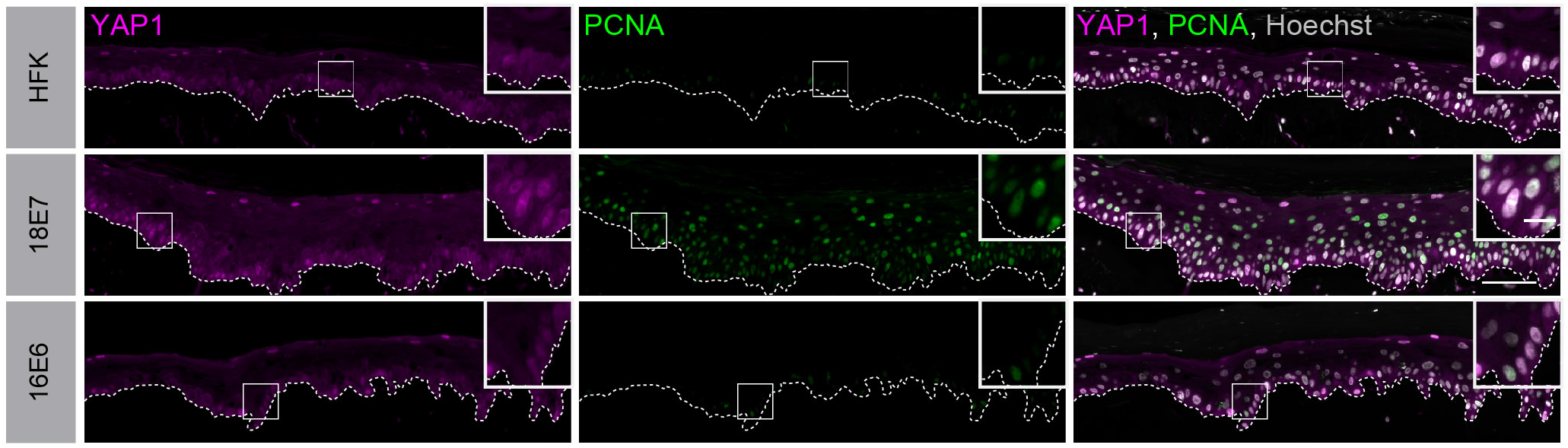
HPV E6 does not activate YAP1 in basal keratinocytes. Additional replicates of organotypic cultures grown from primary HFK or HFK transduced with retroviral expression encoding HPV E6 or E7 proteins. FFPE sections were stained for YAP1 (magenta), PCNA (green), and Hoechst (gray). White dashed lines indicate the basement membrane. White boxes indicate the location of insets in main images. Main image scale bars = 100 μm. Inset scale bars = 25 μm.

**Figure 2—figure supplement 1.**
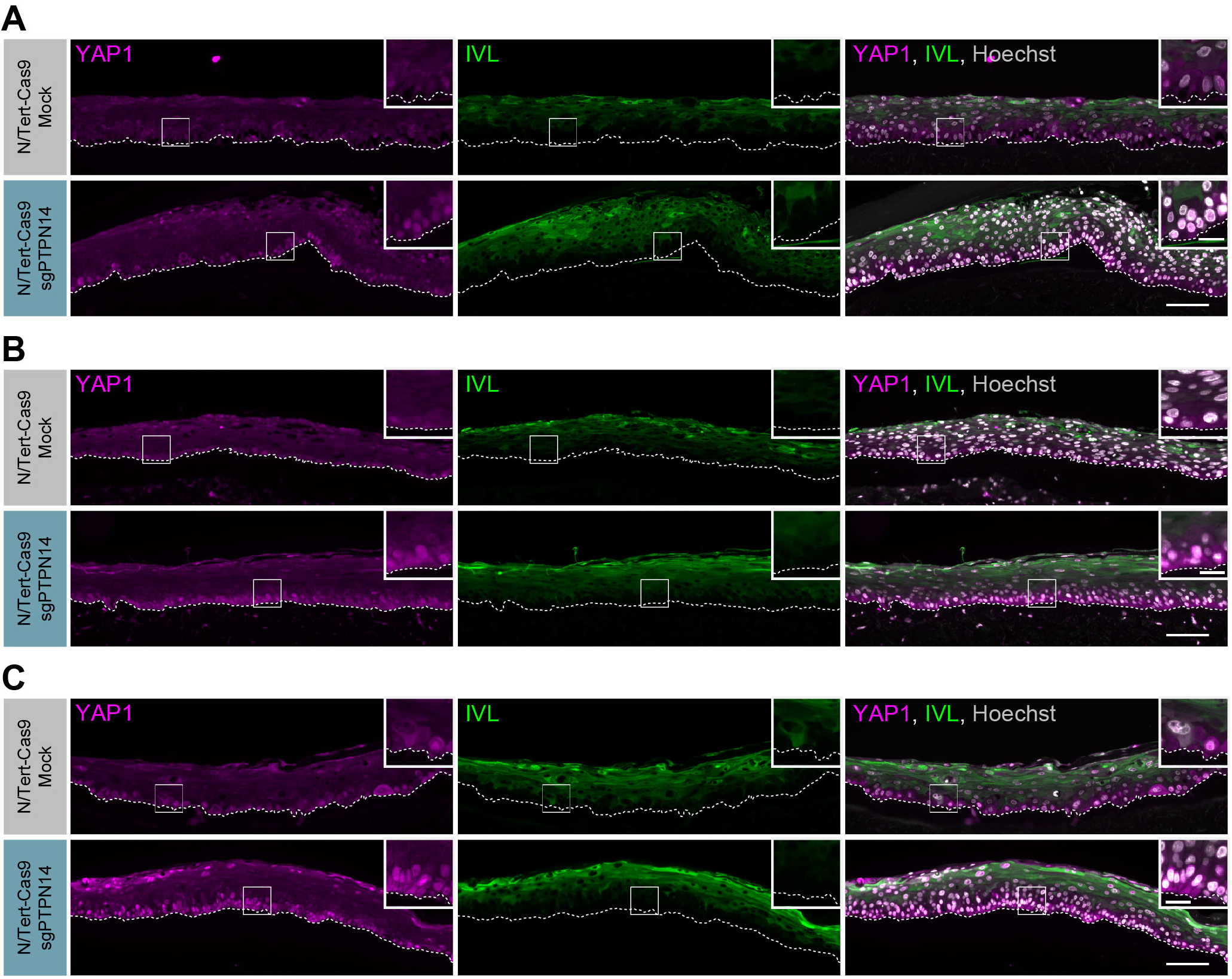
PTPN14 knockout activates YAP1 in basal keratinocytes. Additional replicates of organotypic cultures grown from N/Tert-Cas9 keratinocytes (A-C) FFPE sections from mock or sgPTPN14 transfected N/Tert-Cas9 keratinocytes were stained for YAP1 (magenta), IVL (green), and Hoechst (Gray). White dashed lines indicate the basement membrane. White boxes indicate the location of insets in main images. Main image scale bars = 100 μm. Inset scale bars = 25 μm.

**Figure 2—figure supplement 2.**
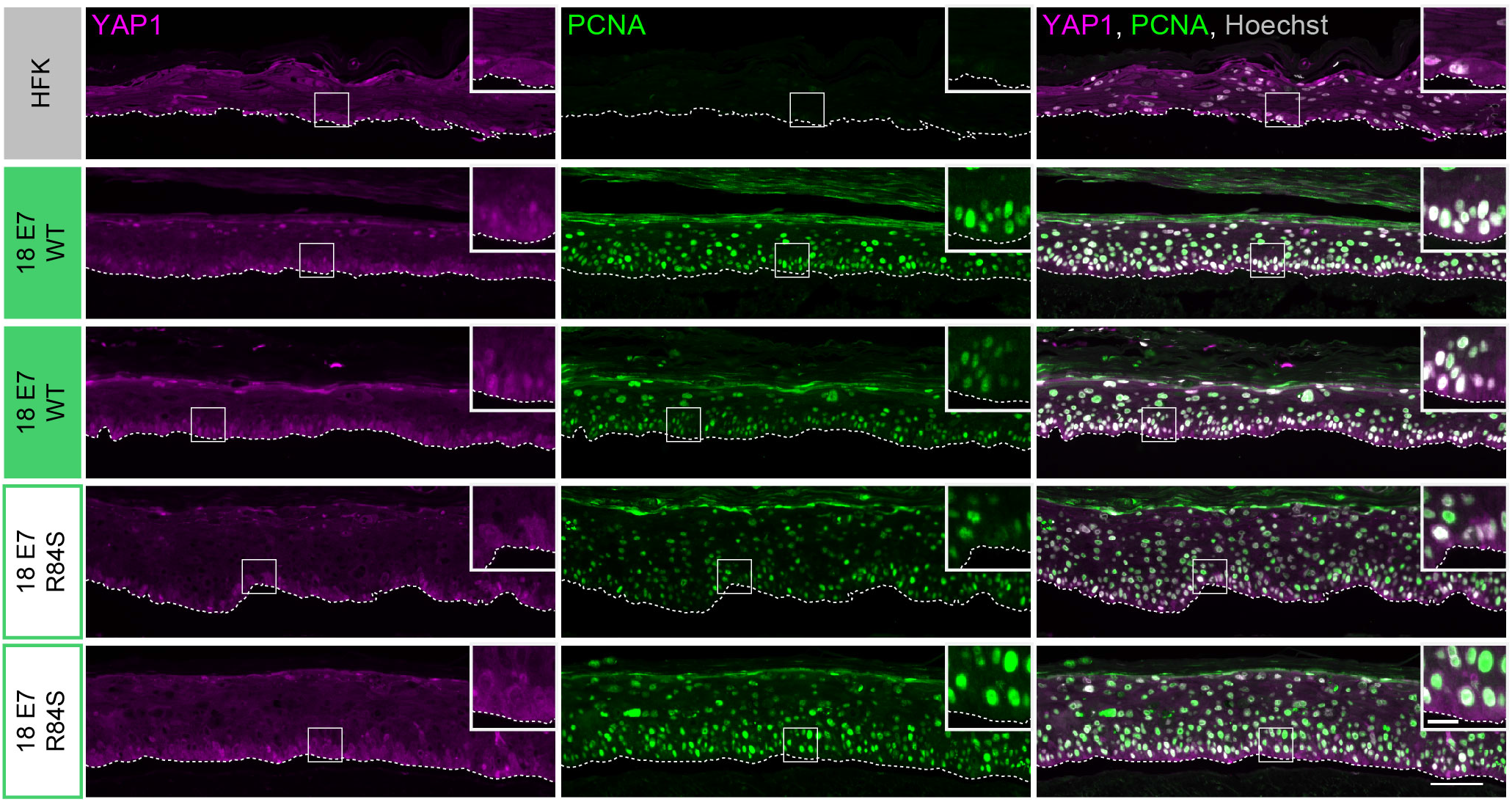
HPV E7 activates YAP1 in basal keratinocytes through PTPN14 degradation. Additional replicates of organotypic cultures grown from primary HFK transduced with retroviral expression vectors encoding HPV18 E7 WT or R84S. FFPE sections from parental HFK, HPV18 E7 WT or HPV18 E7 R84S expressing HFK were stained for YAP1 (magenta), PCNA (green), and Hoechst (Gray). White dashed lines indicate the basement membrane. White boxes indicate the location of insets in main images. Main image scale bars = 100 μm. Inset scale bars = 25 μm.

**Figure 3—figure supplement 1.**
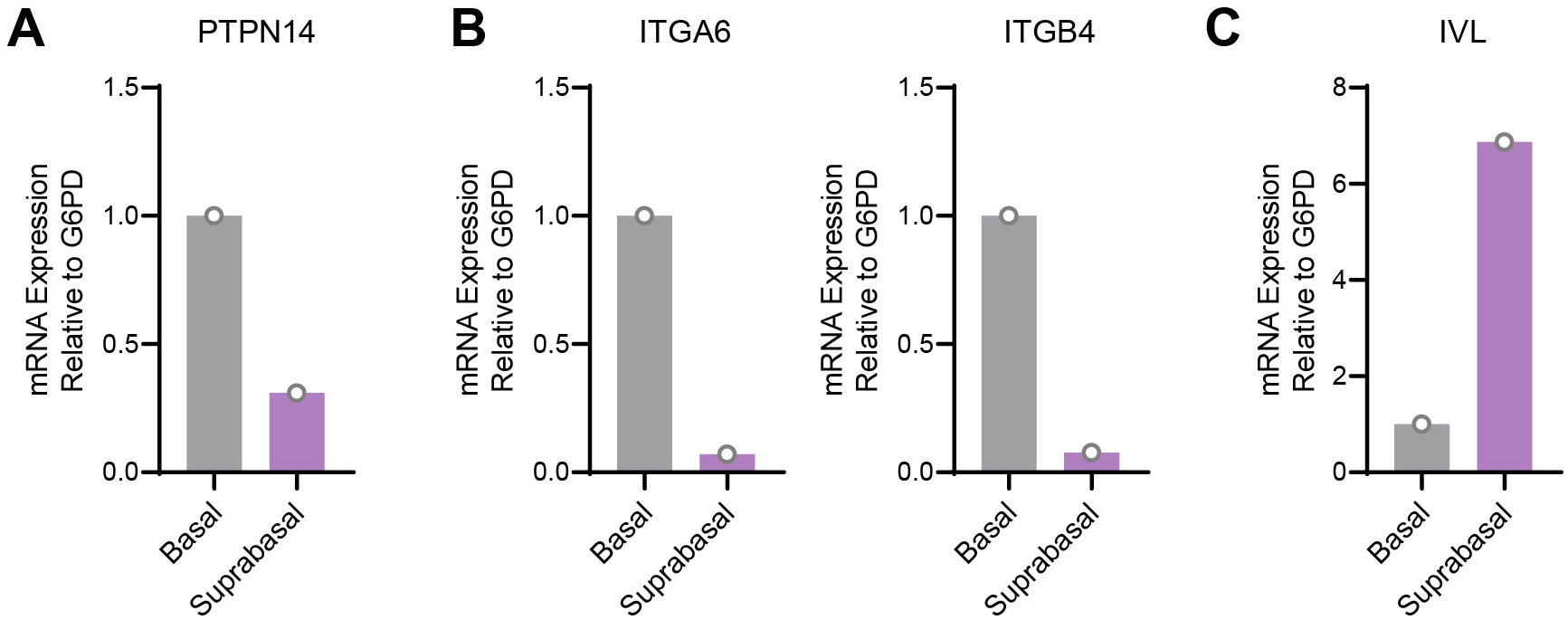
PTPN14 expression is enriched in basal keratinocytes in HPV 18 E7 expressing organotypic cultures. Basal and suprabasal layers from a 3D organotypic culture grown from HFK transduced with a retroviral expression vector encoding HPV18 E7 were dissected using laser capture microdissection. RNA was purified from isolated layers and qRT-PCR was used to assess the expression of PTPN14 (A), the basal cell markers ITGA6 and ITGB4 (B), and the differentiation marker IVL (C). Graphs display individual data points.

**Figure 4—figure supplement 1.**
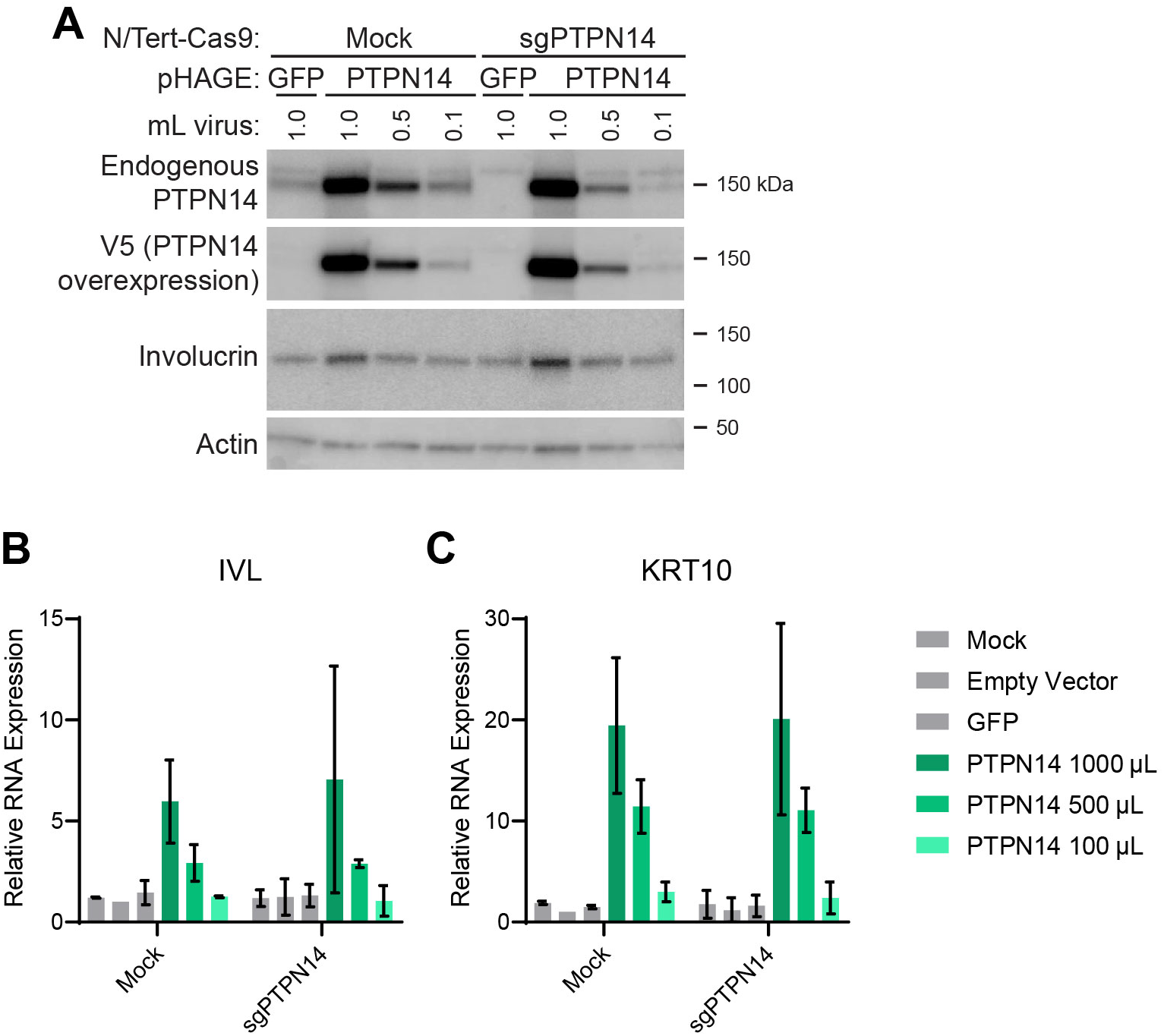
PTPN14 overexpression promotes differentiation in keratinocytes. NTert-Cas9 Mock and sgPTPN14-1 keratinocytes were transduced with lentiviruses encoding GFP or PTPN14 or the empty vector control. (A) Cell lysates were subjected to SDS/PAGE/Western analysis and probed with antibodies to PTPN14, V5-tag, Involucrin, and Actin. (B) qRT-PCR was used to measure the expression of the differentiation markers IVL and KRT10 relative to G6PD. Graphs display the mean ± SD of two independent replicates.

**Figure 4—figure supplement 2.**
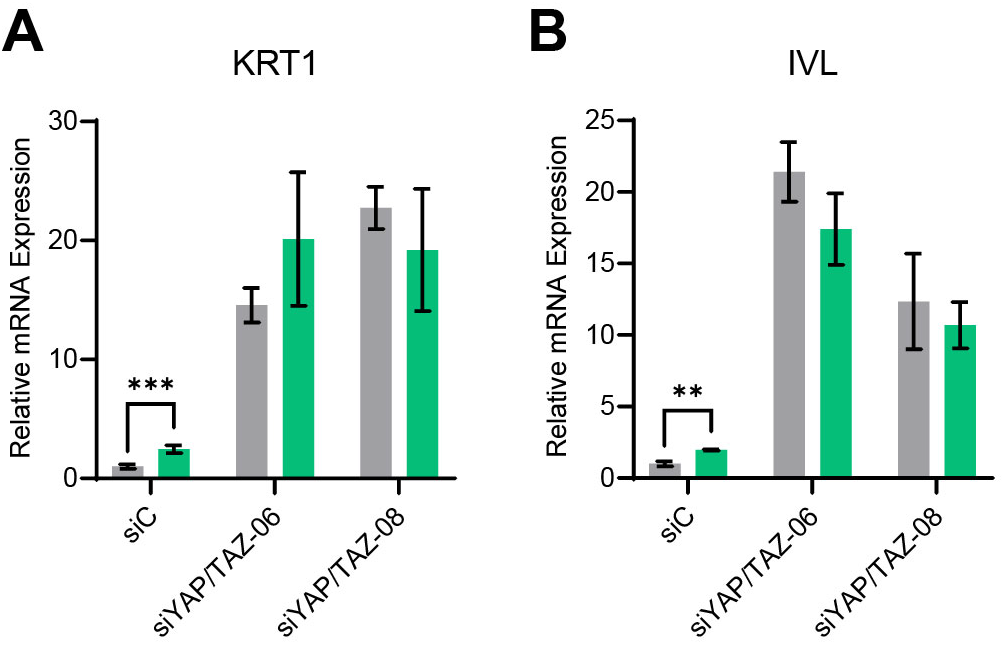
YAP1 and TAZ are required for PTPN14 to promote keratinocyte differentiation. Primary HFK were transfected with control or YAP1 and WWTR1 targeting siRNAs then transduced with PTPN14 encoding lentivirus. qRT-PCR was used to measure the expression of the differentiation markers (A) KRT1 and (B) IVL relative to G6PD. Graphs portray the change in gene expression relative to siC. Graphs display the mean ± SD of three independent replicates. Statistical significance was determined by ANOVA (**p<0.01, ***p<0.001).

**Figure 5—figure supplement 1.**
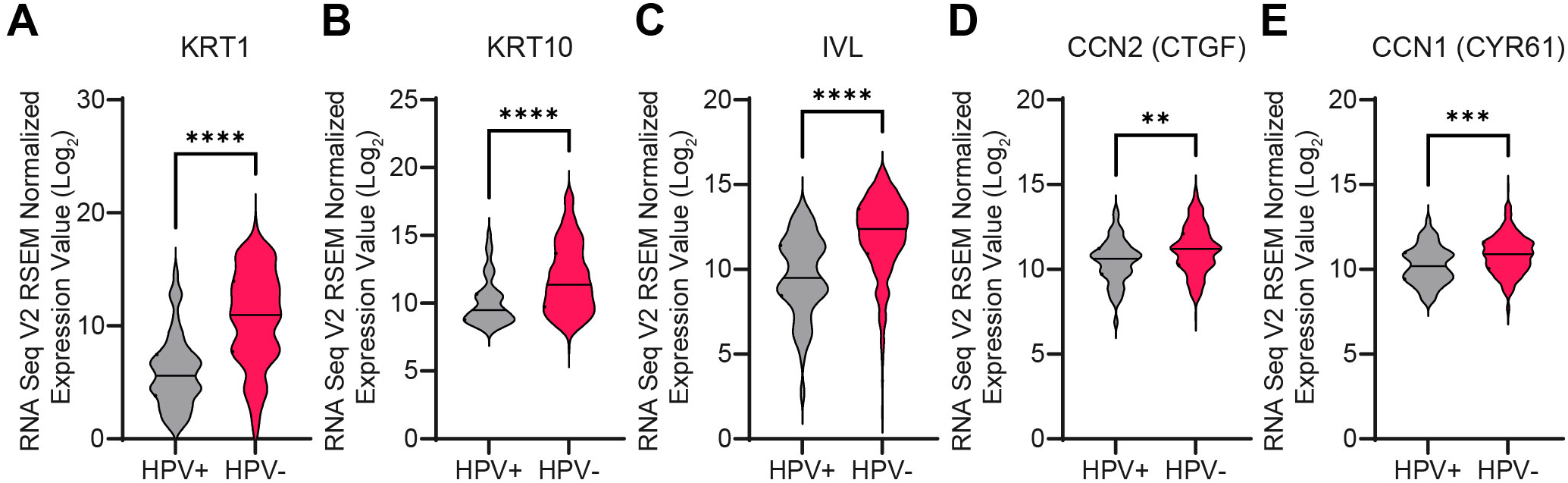
HPV-positive HNSCC express lower levels of differentiation genes. RNA-seq data from TCGA were accessed through cBioPortal. Violin plots display the distribution in log_2_ mRNA expression of differentiation markers (A) KRT1, (B) KRT10, and (C) IVL, and the canonical YAP1/TAZ targets (D) CTGF and (E) CYR61. Statistical significance was determined by Mann-Whitney nonparametric test. (**p<0.01, ***p<0.001, ****p<0.0001).

**Figure 7—figure supplement 1.**
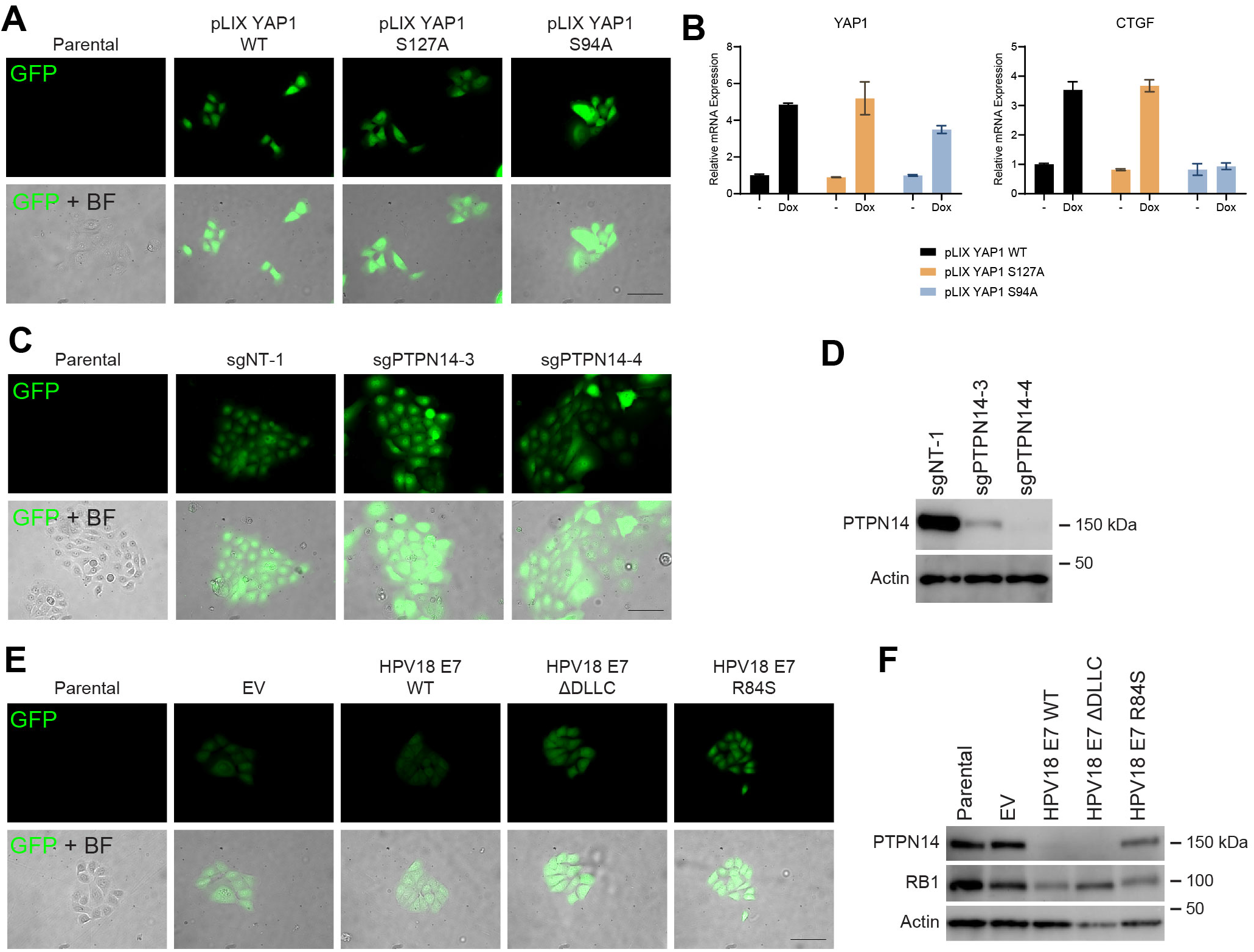
PTPN14 degradation by HPV E7 promotes basal cell retention. (A-B) GFP-labeled HFK were transduced with YAP1 WT, YAP1 S127A, or YAP1 S94A under the control of a doxycycline inducible promoter. (A) GFP expression was confirmed by fluorescence microscopy. Scale bar = 100 μm. (B) Total RNA was purified from monolayer cells +/- treatment with 1 μg/mL doxycycline for 72h. qRT-PCR was used to assess gene expression of YAP1 and CTGF. (C-D) GFP-labeled HFK were transduced with retroviral vectors encoding HPV18 WT, HPV18 ΔDLLC, HPV18 E7 R84S, or the empty vector control (EV). (C) GFP expression was confirmed by fluorescence microscopy. Scale bar = 100 μm. (D) Cell lysates were subjected to SDS/PAGE/Western analysis and probed with antibodies to PTPN14, RB1, and Actin. (E-F) GFP-labeled HFK were transduced with LentiCRISPR v2 sgNT-1, sgPTPN14-3, or sgPTPN14-4 vectors. (E) GFP expression was confirmed by fluorescence microscopy. Scale bar = 100 μm (F) Cell lysates were subjected to SDS/PAGE/Western analysis and probed with antibodies to PTPN14 and Actin.

**Figure 7—figure supplement 2.**
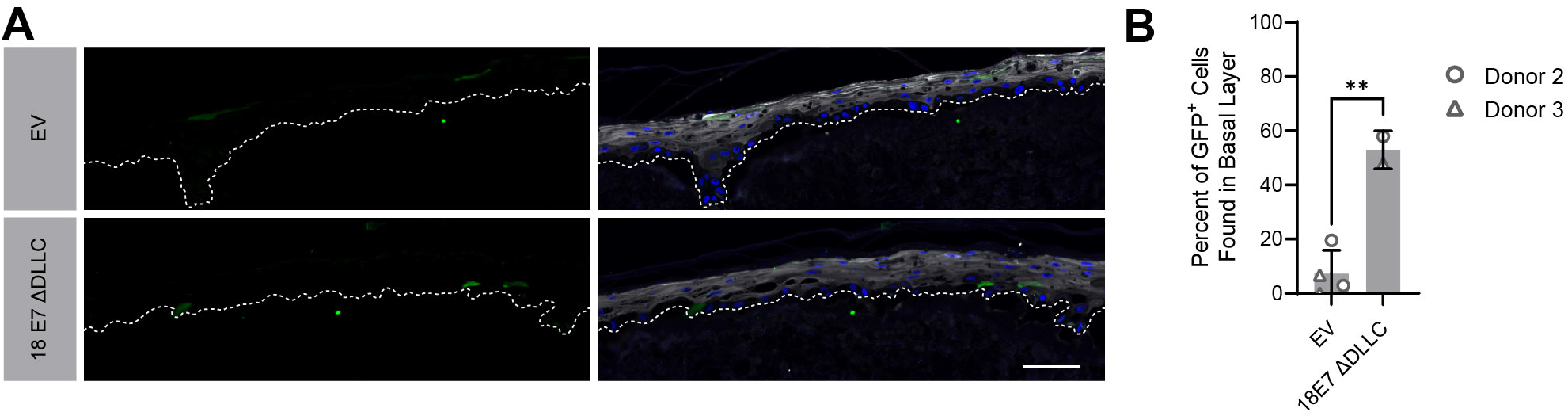
HPV18 E7 can promote basal cell retention in the absence of RB1 binding. Organotypic cultures were grown from GFP-labeled cells mixed with unmodified HFK. GFP-labeled HFK were transduced with HPV18 E7 ΔDLLC or the empty vector (EV). GFP-labeled cells were mixed 1:50 into unmodified HFK. (A) FFPE sections were stained for GFP (green), IVL (grey), and Hoechst (blue). Scale bar = 100 μm (B) Quantification of the percentage of GFP+ cells found in the basal layer. Graphs display the mean ± SD and each individual data point (independent cultures). Statistical significance was determined by t-test. (**p<0.01).

## Key Resources Table

| **Reagent type (species) or resource** | **Designation** | **Source or reference** | **Identifiers** | **Additional information** |
| --- | --- | --- | --- | --- |
| antibody | anti-Actin (Mouse monoclonal) | Sigma-Aldrich | Cat#: MAB1501 | WB (1:20,000) |
| antibody | anti-GFP (Rabbit polyclonal) | Invitrogen | Cat#: A6455 | WB (1:1,000); IHC-P (1:2000) |
| antibody | anti-Mouse IgG Alexa Fluor 488 (Goat polyclonal) | Invitrogen | Cat#: A11001 | IHC-P (1:250) |
| antibody | anti-Mouse IgG HRP (Horse monoclonal) | Cell Signaling Technologies | Cat#: 7076 | WB (1:2000) |
| antibody | anti-Rabbit IgG Alexa Fluor 594 (Goat polyclonal) | Invitrogen | Cat#: A11012 | IHC-P (1:250) |
| antibody | anti-Rabbit IgG HRP (Goat monoclonal) | Cell Signaling Technologies | Cat#: 7074 | WB (1:2000) |
| antibody | anti-HA-Peroxidase (Rat monoclonal) | Roche | Cat#: 12013819001 | WB (1:500) |
| antibody | anti-ITGB4 (Rabbit polyclonal) | Sigma-Aldrich | Cat#: HPA036348 | IHC-P (1:100) |
| antibody | anti-IVL (Mouse monoclonal) | Santa Cruz Biotechnology | Cat#: sc-398952 | IHC-P (1:100) |
| antibody | anti-KRT1 (Mouse monoclonal) | Enzo Life Sciences | Cat#: C34904 |  |
| antibody | anti-PCNA | Santa Cruz Biotechnology | Cat#: sc-56 | IHC-P (1:100) |
| antibody | Anti-PTPN14<br>(Rabbit monoclonal) | Cell Signaling<br>Technology | D5T6Y; Cat#:<br>13808 | WB (1:500) |
| antibody | anti-TAZ<br>(Rabbit monoclonal) | Cell Signaling<br>Technology | D3I6D; Cat#:<br>70148 | WB (1:1000) |
| antibody | anti-V5 (Mouse<br>monoclonal) | Invitrogen | Cat#: 46-0705 | WB (1:1000) |
| antibody | anti-YAP1<br>(Rabbit monoclonal) | Cell Signaling<br>Technology | D8H1X; Cat#:<br>14074 | WB<br>(1:1000);<br>IHC-P (1:50) |
| transfected<br>construct<br>(human) | nontargeting<br>siRNA | Dharmacon | Cat#: D-001810-01 |  |
| transfected<br>construct<br>(human) | siRNA to YAP1<br>(OnTarget Plus) | Dharmacon | Cat#: J-012200-06 |  |
| transfected<br>construct<br>(human) | siRNA to YAP1<br>(OnTarget Plus) | Dharmacon | Cat#: J-012200-08 |  |
| transfected<br>construct<br>(human) | siRNA to<br>WWTR1<br>(OnTarget Plus) | Dharmacon | Cat#: J-016083-06 |  |
| transfected<br>construct<br>(human) | siRNA to<br>WWTR1<br>(OnTarget Plus) | Dharmacon | Cat#: J-016083-08 |  |
| transfected<br>construct<br>(human) | siRNA to<br>PTPN14<br>(OnTarget Plus) | Dharmacon | Cat#: J-008509-05 |  |
| transfected<br>construct<br>(human) | siRNA to<br>PTPN14<br>(OnTarget Plus) | Dharmacon | Cat#: J-008509-08 |  |
| transfected<br>construct<br>(human) | siRNA to<br>LATS1<br>(OnTarget Plus) | Dharmacon | Cat#: J-004632-05 |  |
| transfected<br>construct<br>(human) | siRNA to<br>LATS1<br>(OnTarget Plus) | Dharmacon | Cat#: J-004632-08 |  |
| transfected<br>construct<br>(human) | siRNA to<br>LATS2<br>(OnTarget Plus) | Dharmacon | Cat#: J-003865-09 |  |
| transfected<br>construct<br>(human) | siRNA to<br>LATS2<br>(OnTarget Plus) | Dharmacon | Cat#: J-003865-10 |  |

---

**Supplemental File 1.**
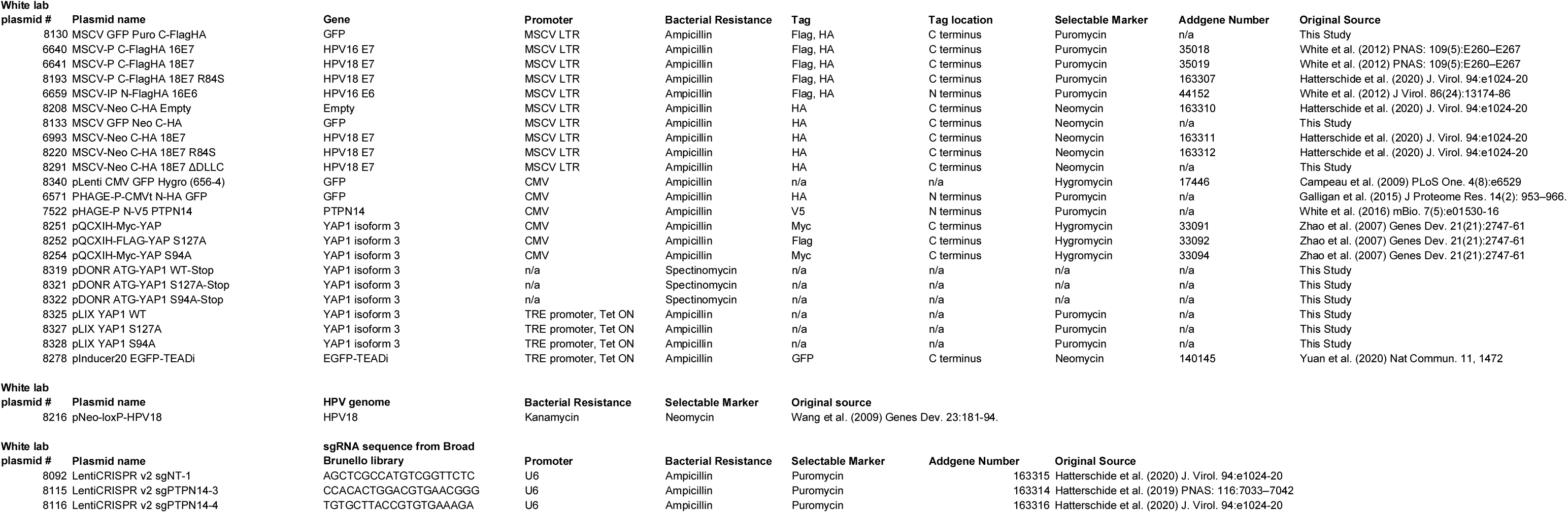
Plasmids used in the study

**Supplemental File 1.**
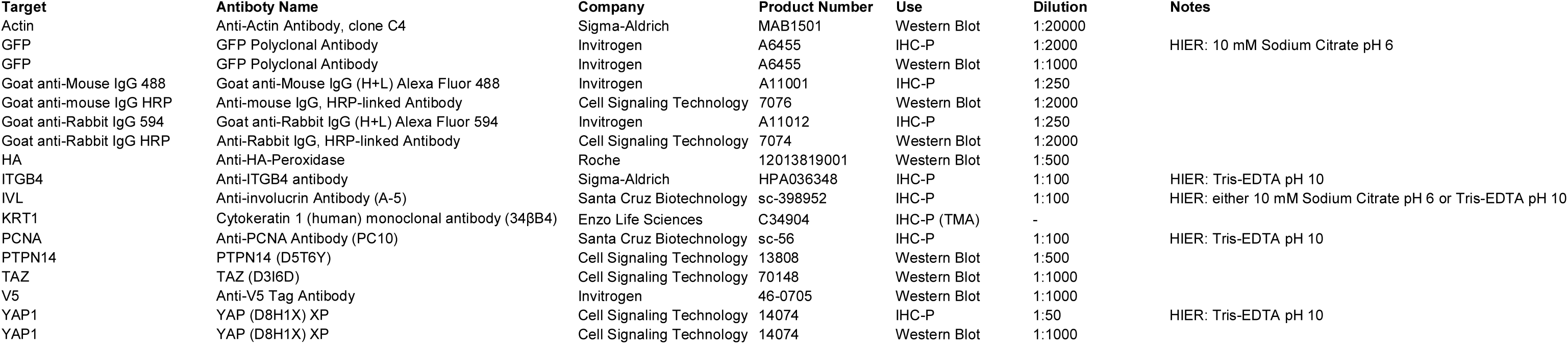
Antibodies used in the study

**Supplemental File 2.**
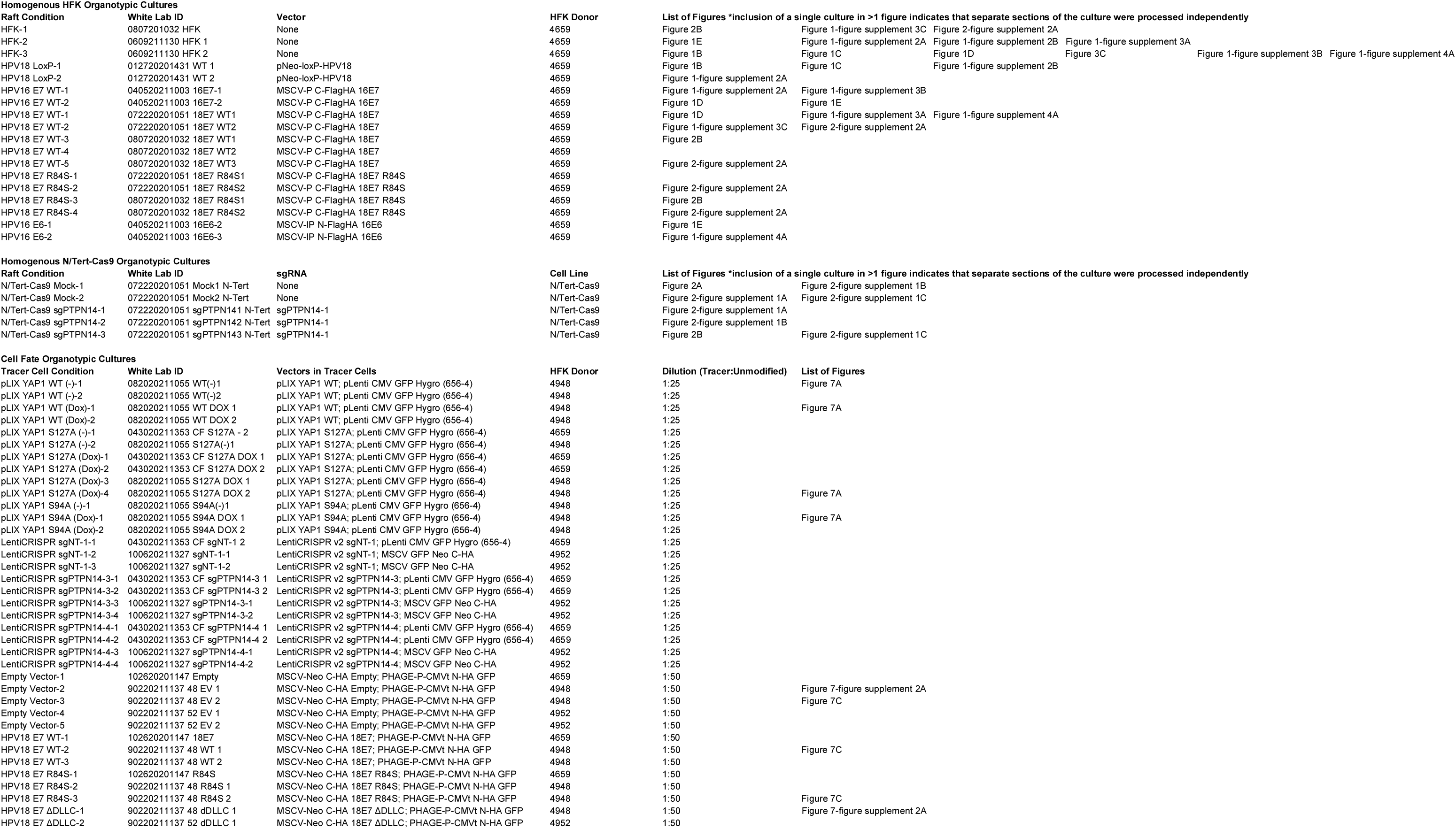
Organotypic cultures used in the study

**Supplemental File 3.** Tumor microarray specimen information

independently **Supplemental File 3****Tumor microarray specimen information**
|  | **Oral Cavity** | **Oropharynx** | **Total** | **HPV-positive** | **HPV-negative\*** |
| --- | --- | --- | --- | --- | --- |
| **# Patients** | 72 | 48 | 120 | 33 | 87 |
| **Primary tumor (T-stage) †** |  |  |  |  |  |
| Early (T1 or T2) | 60 | 40 | 100 | 27 | 73 |
| Advanced (T3 or T4) | 12 | 8 | 20 | 6 | 14 |
| **Nodal metastasis** |  |  |  |  |  |
| Positive | 24 | 39 | 63 | 29 | 34 |
| Negative | 48 | 9 | 57 | 4 | 53 |
| **Overall pathologic stage †** |  |  |  |  |  |
| Early (I or II) | 43 | 7 | 50 | 3 | 47 |
| Advanced (III or IV) | 29 | 41 | 70 | 30 | 40 |

\*HPV status was defined by IHC for p16 for oropharyngeal tumors during routine clinical and was inferred as negative for oral cavity tumors per standards of the College of American Pathologists (Lewis et al. (2018) Archives of Pathology & Laboratory Medicine 142:559–597).
† 7th edition AJCC staging manual

**Supplemental File 4.**
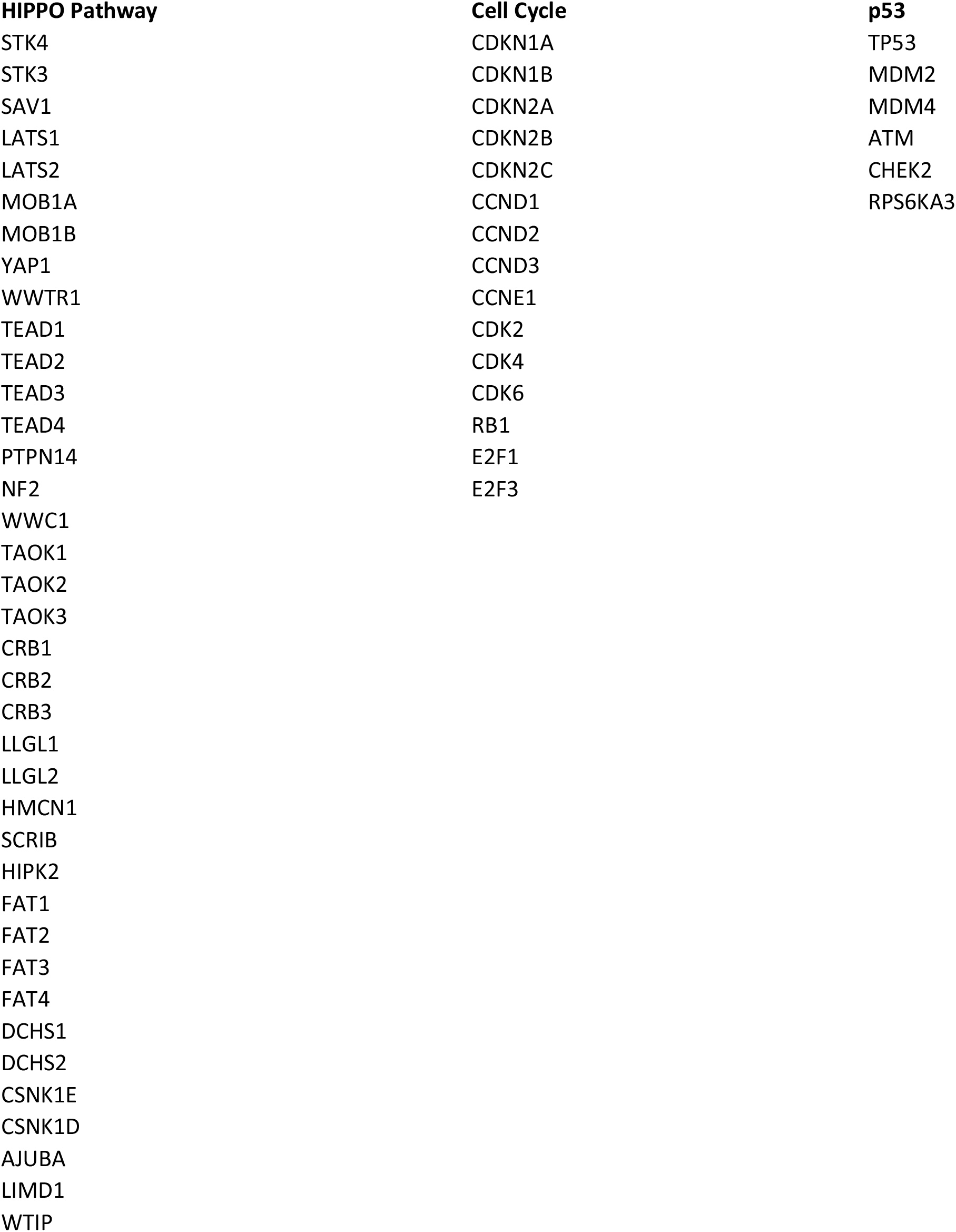
Gene Lists for Pathway Mutational Analyses

